# The Arabidopsis deNADding enzyme DXO1 modulates the plant immunity response

**DOI:** 10.1101/2025.03.11.642587

**Authors:** Anna Golisz-Mocydlarz, Monika Zakrzewska-Placzek, Michał Krzyszton, Nataliia Diachenko, Justyna Piotrowska, Wiktoria Kalbarczyk, Agnieszka Marasek-Ciolakowska, Joanna Kufel

## Abstract

- DXO1, the only DXO homolog in Arabidopsis, due to its plant-specific features, has a strong deNADding enzymatic activity but no apparent role in 5′QC. Most molecular and morphological changes observed so far in *dxo1* mutant plants depended on the plant-specific N-terminal domain of the protein.
- Our investigation revealed the importance of DXO1 enzymatic activity in the plant immune response. We observed that *dxo1-2* knockout mutant and transgenic *dxo1-2* lines expressing a DXO1 variant either catalytically inactive or lacking the N-terminal domain exhibited enhanced resistance to *Pseudomonas syringae*, accompanied by marked changes in the expression of key pathogenesis markers. Other markers of plant immunity, such as callose deposition and production of ROS, were strongly induced by elf18 and flg22.
- Our results strongly suggest that both DXO1 features, the N-terminal domain and its catalytic site, contribute to regulating plant immunity. This is the first observation revealing the involvement of the DXO1 enzymatic activity in plant physiology.
- Moreover, our analyses showed that *dxo1-2* mutation altered the expression of a large group of defense-related genes, affected their mRNAs’ stability, and delayed activation of MAP kinases. Therefore, we postulate that DXO1 protein deregulates defense against *Pst* infection at both the transcriptional and post-transcriptional level.

## Introduction

During their life cycle, plants are exposed to various environmental stresses, many of which are caused by bacteria. The best-characterized gram-negative plant-infecting bacteria is *Pseudomonas syringae* (*Pst*), one of the most agricultural plants pathogens, with more than 50 pathovars (O’Brien *et al*., 2011; Xin *et al*., 2018). *P. syringae* usually enters the leaves through open stomata and colonizes the intercellular space of the apoplast, where it multiplies intensively, leading to visible disease symptoms. The first step of the plant response to infection is the activation of the pattern-triggered immunity (PTI) response pathway, which is induced by pathogen-associated molecular patterns (PAMPs) that bind pattern recognition receptors (PRRs) (Xin & He, 2013; Macho & Zipfel, 2014; Wu *et al*., 2016; Tang *et al*., 2017). The best-known PRRs are a leucine-rich repeat receptor-like kinase (LRR-RLK) FLS2 (FLAGELLIN SENSITIVE 2) and the receptor kinase EFR (ELONGATION FACTOR Tu RECEPTOR), which recognize conserved bacterial epitopes flg22 and elf18, respectively. FLS2/EFR recruit the receptor like-kinase BAK1 (BRI1-ASSOCIATED RECEPTOR KINASE) to form a complex and then phosphorylate the kinase BIK1 (BOTRYTIS-INDUCED KINASE 1), which is released to activate downstream signaling components such as mitogen-activated protein kinase (MAPK) cascades, WRKY transcription factors, reactive oxygen species (ROS) burst, transcriptional reprogramming and phytohormone signaling (salicylic acid (SA), jasmonic acid (JA), abscisic acid (ABA), and ethylene (ET)) (Li *et al*., 2016; Xin *et al*., 2018; Yuan *et al*., 2021). One of the first responses of the PTI is stomatal closure to limit bacterial entry into the apoplast, which is driven by activated SA and ABA signaling pathways (Melotto *et al*., 2008; Cao *et al*., 2011; Lim & Lee, 2015; Xin *et al*., 2018). The second step of the plant response to infection is called effector-triggered immunity (ETI), in which intracellular receptors (NLRs) directly or indirectly recognize pathogen virulence molecules called effectors (Dodds & Rathjen, 2010; Muthamilarasan & Prasad, 2013; Yuan *et al*., 2021). *Pst* delivers effectors into the host cell through the type III secretion system (T3SS), thereby increasing bacterial virulence, but plants can recognize effectors through resistance (R) proteins (Truman *et al*., 2006; Cunnac *et al*., 2009; Xin *et al*., 2018). In general, PTI and ETI cause similar responses, although ETI is more potent and often leads to local cell death.

In response to biotic stress that affects growth and development, plants reprogram cellular metabolism and activate the immune response pathways to adapt to stress conditions. This depends on efficient RNA metabolism, particularly transcription, RNA maturation, and degradation (Kufel *et al*., 2022). One of the RNA-related factors that has recently been reported to have an impact on plant defense response in Arabidopsis is DXO1 (Kwasnik *et al*., 2019; Pan *et al*., 2020).

Eukaryotic DXO/Rai1 family proteins are involved in mRNA 5′-end quality control (5′QC) (Jiao *et al*., 2010; Chang *et al*., 2012) and removal of the non-canonical caps such as NAD^+^ (deNADding), FAD (deFADding) or dpCoA (deCoAping) (Jiao *et al*., 2017; Kiledjian, 2018; Kwasnik *et al*., 2019; Doamekpor *et al*., 2020; Pan *et al*., 2020). Depending on the organism, DXO/Rai1 proteins have different additional enzymatic activities. For example, of the two yeast orthologs, the nuclear Rai1 possesses 5′ pyrophosphatase (PPH) activity and can also remove an unmethylated cap (Xiang *et al*., 2009; Jiao *et al*., 2010), whereas Dxo1 lacks PPH activity but can hydrolyze caps and is also a 5′-3′ exoribonuclease (Jiao *et al*., 2010; Chang *et al*., 2012). In turn, mammalian DXO is a hybrid of fungal Rai1 and Dxo1 and has multiple functions as a PPH, 5′-3′ exoribonuclease, and in decapping of non-canonical caps (Jiao *et al*., 2013, 2017; Grudzien-Nogalska & Kiledjian, 2016).

*Arabidopsis thaliana* DXO1 protein, like its homologs in other plant species, shares a high degree of active site conservation with its mammalian and fungal counterparts. In addition, Arabidopsis DXO1 have a plant-specific 194 amino acid N-terminal extension (NTE) that stabilizes protein-RNA interactions, and an amino acid substitution near the active site that affects its enzymatic properties (Kwasnik *et al*., 2019). As a result, DXO1 possesses strong deNADding and weak 5′-3′ exoribonuclease activities but almost no 5′QC activity (Kwasnik *et al*., 2019; Pan *et al*., 2020). It has recently been reported that DXO1 is also an essential component of the m^7^G capping machinery by activating the RNA guanosine-7 methyltransferase RNMT1, which catalyzes methylation of the guanosine cap (Xiao *et al*., 2023).

DXO1 deficiency results in growth retardation, developmental defects and pale-green coloration, altered response to biotic and abiotic stress, as well as molecular changes, namely deregulation of expression of specific classes of genes, ineffective mRNA capping, accumulation of RDR6-dependent siRNAs, and defects in rRNA maturation (Kwasnik *et al*., 2019; Zakrzewska-Placzek *et al*., 2025). Intriguingly, these phenotypes of *dxo1* plants and most DXO1 cellular functions, except the recently described mechanism of co-translational mRNA decay (CTRD) (Carpentier *et al*., 2025), do not require its catalytic activity but the plant-specific N-terminal extension.

DXO1 has been shown to be involved in modulating the response to different stresses. Its presence is important for proper ABA signaling during seed germination, reaction to salt, high temperature, drought, wounding, and sucrose or glucose starvation (Pan *et al*., 2020; Yu *et al*., 2021; Zakrzewska-Placzek *et al*., 2025). In addition, the *dxo1* mutation causes constitutive activation of the immune pathway and increased resistance to infection by *Pseudomonas syringae* (Pan *et al*., 2020). At the molecular level, the lack of DXO1 results in a strong upregulation of defense-related genes such as *PR1*, *PR2*, and *PAD4*, and downregulation of photosynthesis-related genes (Kwasnik *et al*., 2019; Pan *et al*., 2020).

In this work, we analyzed in detail the role of Arabidopsis DXO1 protein in response to biotic stress induced by *Pseudomonas syringae* pv. *tomato* DC3000. We confirmed that the DXO1-deficient *dxo1-2* mutant is resistant to *Pst*, but we also observed an unusual biphasic response in which plants were initially hypersensitive to the pathogen and only later became resistant. Our genome-wide transcriptome analysis of the *Pst*-infected *dxo1-2* mutant revealed that the lack of DXO1 deregulates defense against *Pst* infection at the transcriptional level. In turn, the impact of *dxo1-2* mutation on the stability of mRNAs encoding pathogen-related factors suggests that this protein contributes to the regulation of plant immunity also at the posttranscriptional level. Finally, we tested transgenic *dxo1-2* lines expressing different DXO1 variants, wild-type (DXO1(WT)), catalytic mutant (DXO1(E394A/D396A)), and lacking the N-terminal domain (DXO1(ΔN194)), and found that they also showed increased resistance to *Pst* and marked changes in the expression of major pathogenic markers. Other indicators of plant resistance, such as elf18- and flg22-induced callose deposition, production of reactive oxygen species and activation of MAPKs, were also altered in all tested lines. These results indicate that, in contrast to other phenotypes described to date, the response to biotic stress induced by *Pst* depends not only on the N-terminal domain of DXO1 but also on its catalytic activity. Interestingly, specific differences between these lines indicate that NTE- and enzymatic-mediated response pathways are at least partially distinct.

## Materials and Methods

### Plant material and growth conditions

*Arabidopsis thaliana* wild-type ecotype Columbia (Col-0) and mutant *dxo1-2* (SALK_032903) plants were used in this study. We also used transgenic *dxo1-2* lines expressing different DXO1 variants: DXO1(WT), DXO1(E394A/D396A), and DXO1(ΔN194) fused to GFP under the control of the constitutive 35S promoter from the cauliflower mosaic virus (CaMV) (Kwasnik *et al*., 2019). Seeds were surface sterilized with 30% bleach and 0.02% Triton-X100 solution and sown on MS medium (Murashige & Skoog, 1962) supplemented with 1% (w/v) sucrose and 0.3% phytagel. After 3 days of stratification at 4°C, plates were moved to growth chambers under a 16 h light/8 h dark (long-day) photoperiod at 22/19°C and seedlings were harvested after 2 weeks. Infection experiments were performed on 6-week-old plants grown in soil under an 8 h light/16 h dark (short-day) photoperiod at 22/19°C.

### Bacterial infection assays and PAMP treatments

Bacterial infection assays with the virulent *Pseudomonas syringae* pv. *tomato* strain DC3000 (*Pst*) was performed as described previously (Golisz *et al*., 2021). Briefly, 6-week-old plants were inoculated by spraying or injection with the *Pst* suspension in 10 mM MgCl_2_. Material was harvested from at least 10 plants for each time point, frozen in liquid nitrogen and used for RNA extraction. Bacterial growth was quantified by assessing the number of dividing bacterial cells 24 h, 48 h and 72 h after infection (hpi). Samples (four leaf discs) were taken using a cork-borer (4 mm) from 2 leaves, ground in sterile Mili-Q water, diluted and plated on selection medium. Plates were incubated at 28°C, and colonies were counted after 48 h.

PAMP assays were performed as described previously (Golisz *et al*., 2021). In brief, 14-day-old seedlings were treated with flg22 (Alpha Diagnostic International Inc.) and elf18 (synthesized by GL Biochem Ltd, Shanghai, China) to a final concentration of 100 nM. Seedlings were harvested at the indicated time points and frozen in liquid nitrogen.

### RNA methods

Total RNA was isolated from 14-day-old seedlings or 6-week-old plants using TRI Reagent (Sigma-Aldrich) according to the manufacturer’s instructions. High-molecular-weight RNAs were analysed in 1.1% agarose gel and transferred to a Hybond N+ membrane by capillary elution. Northern blots were carried out using α-ATP32 random primed probes and the DECAprimeTM II labelling kit (ThermoFisher Scientific). Quantification of northern blots was performed using a typhoon FLA 9000 Gel Imaging Scanner (GE Healthcare) and ImageQuant software (Molecular Dynamics).

mRNA half-life measurement experiments were carried out as described (Kwasnik *et al*., 2019). Two-week-old seedlings were transferred to flasks containing a buffer (1 mM PIPES, pH 6.25, 1 mM sodium citrate, 1 mM KCl, 15 mM sucrose), and after a 30-min incubation, cordycepin (150 mg/l) was added. Total RNA samples corresponding to 0, 15, 30, 60, 90 and 120 min time points after transcriptional inhibition were extracted using TRI Reagent (Sigma-Aldrich) and analysed by northern blot. Oligonucleotides used for northern hybridization are listed in Supplementary Table S1.

### 3′RNA-seq

We performed RNA sequencing according to recently published work (Krzyszton *et al*., 2022). RNA was isolated as described above from mock or pathogen-treated Col-0 and *dxo1-2* plants in four replicates. After Turbo DNase treatment, 500 ng of RNA was reverse transcribed using SuperScript III and barcoded oligo (dT) primers (Supplementary Table S1) in 20 µl reactions. 5 µl of each reaction were then pooled, and combined cDNA was concentrated with Ampure RNA Clean-up to 15 µl. The recovered cDNA was mixed with 15 µl of second-strand synthesis mix (2x NEBNext Second Strand Synthesis (dNTP-free) Reaction Buffer (NEB), 1 U RNaseH (NEB), 1 U E. coli DNA ligase I (NEB), 5 U E. coli DNA polymerase (NEB), 30 µM dNTPs) and incubated overnight at 16°C. Double-stranded DNA was purified with Ampure beads (1.2x beads to sample volume) and eluted in 3 µl of water. Tagmentation was performed with homemade Tn5 (3 µl) and 6 µl freshly prepared 2x buffer (20 mM Tris-HCl pH 7.5, 20 mM MgCl2, 50% DMF). Tn5 enzyme was purified according to (Hennig et al., 2018), loaded with B duplex and diluted five times before use. The reaction was mixed and incubated for 7 min at 55°C followed by 5 min at 80°C. Water was added up to 30 µl and samples were purified with Ampure beads (1.2x beads to sample volumes). Illumina indexing PCR was performed using 20 µl tagmented DNA in 50 µl reactions with Q5 2× PCR Master Mix (NEB) and 0.5 µM final concentration of Nextera indexing primers. To avoid PCR over-cycling, we estimated the number of cycles as in (Buenrostro et al., 2015). Libraries were sequenced on the Illumina NextSeq 500 system using the paired-end mode to obtain 21 nt R1 (containing barcode and UMI) and 55 nt R2 (containing mRNA sequence).

Processing of the data: In our oligo (dT) primers, two parts of UMI are split by barcode sequence, which is why we transformed read R1 fastq file using awk command: *awk ‘NR%2==1 {print $0} NR%2==0 {print substr($1,16,5) substr($1,1,15)}’*. Read R2 was trimmed to remove potential contamination with poly(A) tail using BRBseqTools (v 1.6) Trim (Alpern *et al*., 2019) and parameters *-polyA 10 -minLength 30*. Then mapped using STAR (v 2.7.8a) (Dobin *et al*., 2013) to TAIR 10 genome version and Araport11 genome annotation with parameters *--sjdbOverhang 54 --outSAMtype BAM SortedByCoordinate --outFilterMultimapNmax 1*. Finally, the count matrix for each RNA sample and each gene was obtained using BRBseqTools (v 1.6) CreateDGEMatrix (Alpern *et al*., 2019) with parameters *-p UB -UMI 14 -s yes*, using Araport11 genome annotation and a list of barcodes. Counts were used for differential gene expression analysis and PCA using the DESeq2 R package (Love *et al*., 2014). Overlaps between gene sets were visualised with UpSetR (Conway *et al*., 2017) and GO-terms enrichments were calculated with gprofiler2 (Kolberg *et al*., 2020) using expressed genes as a background. For gene expression clustering, we used the WGCNA R package (Langfelder & Horvath, 2008).

In this work, we reanalysed 3’RNA-seq data for different complementation lines of *dxo1-2* mutant using gene expression counts submitted to GEO under GSE210631 accession number. Measurement of apoplastic ROS production Flg22-triggered H_2_O_2_ production was detected using GloMax®-Multi^+^ Detection System (Promega) according to published protocols with minor modifications (Smith & Heese, 2014; Bisceglia *et al*., 2015). Leaf disks cut from 6-week-old plants were submerged adaxial side up in 200 μl of water in individual wells of a 96-well microplate for 24 hours at 22°C to reduce the wounding response. Immediately prior to measurement, water was gently replaced with an equal volume of luminol/peroxidase. One of the injectors was charged with the 5× flg22 solution. The luminescence detection assays were performed for at least 30 min with 1 sec signal integration time. The luminescence was in relative light units (RLU).

### Callose deposition assay

Ten sterilized Col-0 and *dxo1-2* seeds were sown per well in 6-well plates containing MS medium and grown under long-day conditions for 7 days. The medium was then replaced with fresh MS, and plants were treated with flg22, elf18, and coronatine at the final concentration of 1 mM for 24 h, as described (Luna *et al*., 2011). Samples were washed with 95% EtOH and incubated for 2 h in 0.07 M K_2_HPO_4_ containing 0.01% aniline blue (Sigma). Imaging was performed using a fluorescence microscope with a DAPI filter at a wavelength of 370 nm, and images were analyzed using the ImageJ software.

### Leaf phenotypical traits

Stomatal characteristics: The abaxial epidermis was isolated from the middle of five leaves of each genotype using transparent adhesive tape and stained with 2% toluidine blue. The length of stomata (n = 5 replication × 30 stomata/genotype) and their density per 1 mm^2^ (n = 5 replication/genotype) were determined using a Nikon Eclipse 80i microscope (Nikon) at 100 and 400 times magnification employing an image analysis system NIS-Elements BR ver. 2.30 (Nikon Instruments Inc.).

Trichome density: Fresh leaves that were not fixed were used for trichome morphology and density analyses. Images of the abaxial and adaxial leaf surfaces were captured using a digital microscope (VHX-7000N Keyence) at 40x magnification. Four leaves were analyzed for each genotype.

Leaf anatomy: For anatomical observation, 10 × 5 mm pieces of the leaves were excised for each genotype. The material was fixed in chromoacetoformalin (CrAF) solution for 48 h at room temperature, dehydrated through an increasing alcohol series (70, 80, 90, and 100%), and embedded in paraffin according to a previously reported method (Marasek-Ciolakowska *et al*., 2020). Transverse sections (12 μm thick) were cut with a rotary microtome (Leica) and stained with safranin and fast green. The sections were mounted in Canada balsam and analyzed using a Nikon Eclipse 80i microscope (Nikon). For each leaf sample, the thickness of the lamina and thickness of the abaxial and adaxial epidermis were measured at 200 × magnification. For statistical analysis, three replicates were used for each genotype, and each replicate consisted of 30 measurements (n = 3 replicates × 30).

### SA and JA measurements

Homogenized 14-day-old plant tissue was extracted with methanol containing 10 ng mL^-1^ of stable isotope-labeled internal standards: ^2^H_6_-SA, ^2^H_6_-ABA (OlChemIn s.r.o.). The incubation was performed in the safe-lock tubes using a shaker (Eppendorf ThermoMixer) with an acceleration of 1000 rpm at 4°C for 15 min. After keeping the samples in the ultrasonic bath for 5 min, they were centrifuged at 15,000 × g at 4°C for 15 min (Universal 320R, Hettich). The supernatants were transferred into new tubes, concentrated using the Speed-vac system, and then re-suspended with 5 % methanol (v/v). The phytohormone fraction was eluted using acetonitrile and concentrated using the Speed-vac system. The dry pellet was reconstructed in 0.2 mL methanol.

Analysis was performed using the Waters ACQUITY UPLC System (Milford), which is equipped with the Binary Solvent Manager, Sample Manager, and Column Manager. The gradient program of A: 0.1 % formic acid in water (LC-MS grade, Merck-Millipore) and B: 0.1 % formic acid in acetonitrile (LC-MS grade, Merck-Millipore) at a flow rate of 0.15 mL min^-1^ was applied for sample separation on the HSS T3 column (1 × 100 mm, particle size 1.8 μm, Waters) at 40°C. The gradient was started for 1 min at 10 % eluent B and then increased to 50 % and 99 % eluent B for 8 and 10 min, respectively. The column was washed for 3 min at 99 % eluent B, and re-equilibrated for 3 min. Compounds were detected by HESI-HRMS/MS using the QExactive mass spectrometer (Thermo Scientific, Waltham) operated in negative ionization in the parallel reaction monitoring (PRM) and select ion monitoring (SIM) mode.

### MAPK activation assay, protein extraction and western blotting

To detect MAPK phosphorylation, 2-week-old seedlings grown in liquid MS were treated with 100 nM flg22 or elf18 for indicated time points, ground in liquid nitrogen and 300 µl of protein extraction buffer (50 mM Tris-HCl pH 7.5, 100 mM NaCl, 10% glycerol, 2 mM EDTA, 5 mM dithiothreitol, 1% IGEPAL CA630, 2 mM sodium molybdate, 1 mM sodium fluoride, 1 mM sodium orthovanadate, 4 mM sodium tartrate and protease inhibitor cocktail) was added. The suspension was centrifuged at 13 000 *g* for 15 min at 4°C, and the supernatant was collected. Protein concentration in extracts was measured using The Direct Detect® Spectrometer (Merck Millipore). Proteins (50 µg) were separated in 10% SDS-PAGE gels and transferred onto nitrocellulose membrane (GE Healthcare) by electroblotting. Activated MAP kinases were detected with anti-phospho-p42/44 MAPK primary antibodies (1:1000, Cell Signal Technology), followed by incubation with anti-rabbit-HRP secondary antibodies (1:25000, Thermo Scientific). Images were documented using Amersham ImageQuant™ 800 Western blot imaging system (Cytiva).

## Results

### DXO1 deficiency affects the response to *Pst* DC3000 infection

The lack of DXO1 has been reported to confer pleiotropic phenotypes, including altered response to stress (Kwasnik *et al*., 2019; Pan *et al*., 2020; Yu *et al*., 2021; Xiao *et al*., 2023). In particular, the *dxo1-2* knockout mutant showed increased resistance to *Pst* DC3000, constitutive activation of the defense pathway and altered expression of some genes related to plantlpathogen interactions (Pan *et al*., 2020). To look more closely at the role of DXO1 in plant innate immunity, we tested the resistance of the Arabidopsis *dxo1-2* T-DNA insertion mutant to *Pst* DC3000 infection performed by spraying and injection.

Bacterial growth assayed over time after spraying showed that initially, at 24 hpi (hours post-infection), the *dxo1-2* mutant was more sensitive to *Pst* and acquired resistance only later, which is clearly visible at 72 hpi (Fig. 1a). The latter outcome is consistent with the results of previous studies in which *dxo1-2* was resistant to *Pst* three days after infection (Pan *et al*., 2020). To test whether the different early response of the mutant is caused by more effective pathogen penetration, we examined the outcome of plant infection by syringe infiltration, in which bacteria are not delivered into the leaf tissue *via* stomata but directly to the apoplastic space. In the case of injection, the *dxo1-2* plants became more resistant to the pathogen already after 48 hpi (Fig. 1b). The earlier response to pathogen administered by surface inoculation by spraying than by injection suggests that the initial sensitivity of the *dxo1-2* mutant may reflect facilitated pathogen entry at infection sites, possibly via stomatal apertures. We therefore examined the thickness of leaves, the density and size of stomata and the number of leaf hairs, because these traits constitute physical barriers that regulate the entry of bacteria into the plant. The *dxo1-2* mutant has significantly thinner leaves, lower stomatal density and reduced length of stomatal apertures but significantly higher trichome density (Fig. S1, S2a,b). The altered leaf morphology may at least to some extent explain the initially increased susceptibility of *dxo1-2* to infection due to increased *Pst* penetration into the leaf (Mansvelt & Hattingh, 1987; Underwood, 2012; Kim, 2019).

**Fig. 1.**
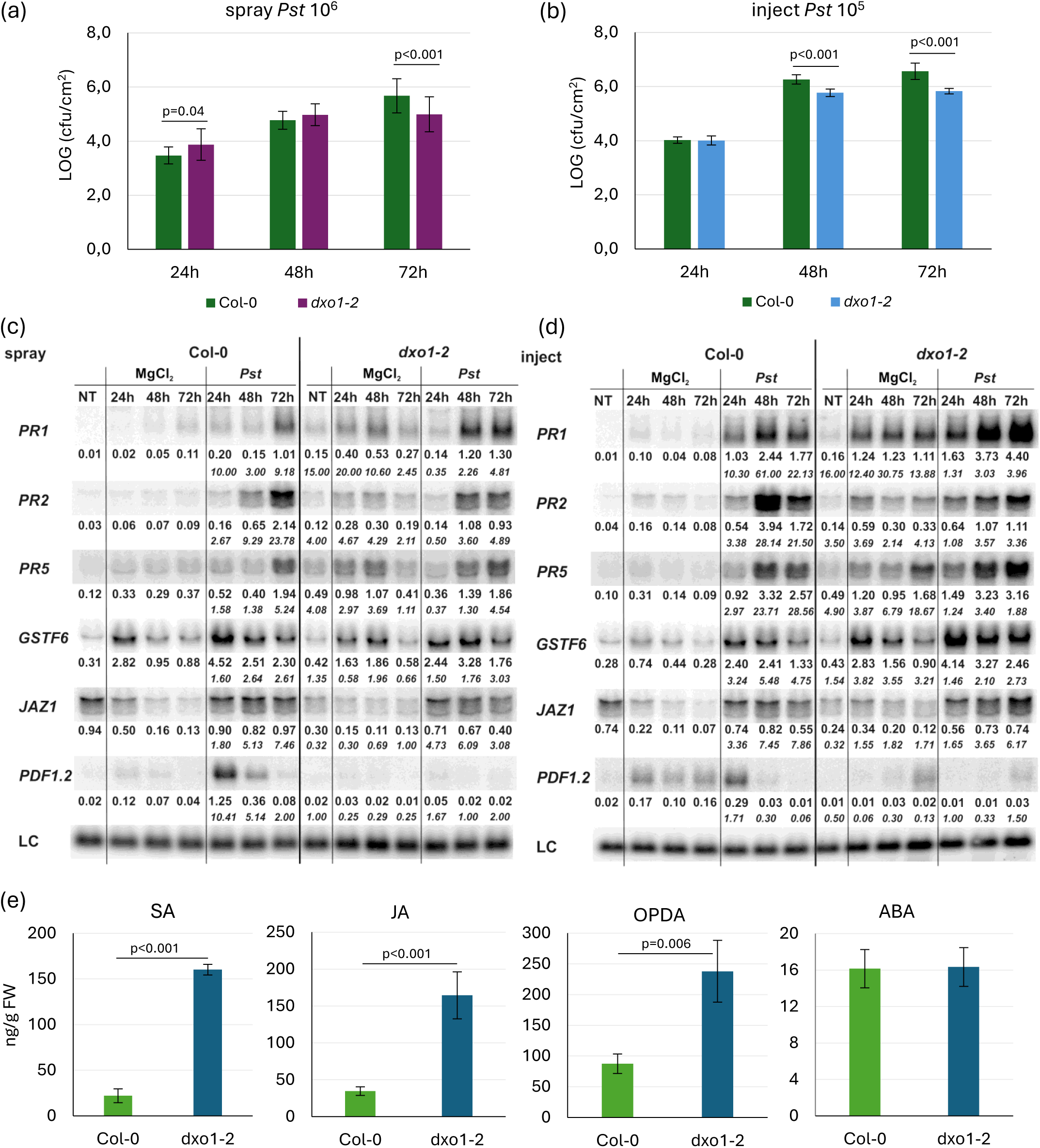
The *dxo1-2* mutant is resistant to *Pseudomonas syringae* pv. *tomato* DC3000. (a, b) Growth of *Pst* DC3000 after 24, 48, and 72 h in Col-0 and the *dxo1-2* mutant after spraying (a) or injection (b). For each time point, leaf discs were collected from 6 plants. Results are the mean of three independent experiments, and error bars represent SD; P < 0.05 (Tukey’s t-test). (c, d) Northern blot analysis of factors involved in response to *Pst* DC3000. Samples were collected from non-treated (NT), control (MgCl_2_) and infected (*Pst*) Col-0 and the *dxo1-2* plants at indicated time points after spraying (c) or injection (d). Numbers represent the ratio of transcript level in treated *dxo1-2* versus Col-0 normalized to U2 snRNA loading control (LC), which is shown as the main numbers, while the ratio relative to the control conditions is given in italics. Experiments were repeated at least three times; representative blots are shown. (e) SA, JA, OPDA, and ABA levels in 14-day-old Col-0 and *dxo1-2* plants. Bars represent the mean of three independent biological replicates with error bars showing SD, P < 0.05 for Tukey’s test.

Next, we examined pathogen-induced changes in the expression of key pathogenesis markers, including *PATHOGENESIS-RELATED GENES* (*PR1*, *PR5*, *PR2*), which are involved in the SA response, *GLUTATHIONE S-TRANSFERASE 6* (*GSTF6*), a marker of ROS oxidative burst, and *PLANT DEFENSIN* (*PDF1.2*) and *JASMONATE-ZIM-DOMAIN PROTEINS* (*JAZ1*) from the JA pathway (Lieberherr *et al*., 2003; Demianski *et al*., 2011; Barah *et al*., 2013; Han *et al*., 2023). Northern blot analysis confirmed the activation of these markers by *Pst* delivered by both infection methods compared to controls (Fig. 1c,d). In the *dxo1-2* mutant, we observed a clear increase in the expression of *PR1*, *PR5*, *GSTF6*, and *JAZ1* after pathogen treatment, especially pronounced in injection-infected plants. In contrast, the expression of *PDF1.2* was strongly downregulated in the *dxo1-2* mutant (Fig. 1c,d). Notably, as previously reported, *PR* genes were visibly activated also in untreated *dxo1-2* plants, consistent with the constitutive defense response phenotype of the mutant (Pan *et al*., 2020). This phenotype is further confirmed by the strongly elevated levels of two plant hormones strictly related to plant immune response, salicylic acid (SA) and jasmonic acid (JA), in the untreated *dxo1-2* line compared to Col-0 (Fig. 1e). The SA pathway plays a major defensive role in response to biotic stress induced by hemibiotrophic pathogens such as *P. syringae*, while the JA pathway primarily affects the response induced by necrotrophic pathogens (Robert-Seilaniantz *et al*., 2011; Pieterse *et al*., 2012). These two defense pathways are mutually antagonistic, but synergistic interactions have also been reported (Kunkel & Brooks, 2002; Derksen *et al*., 2013). In particular, high basal levels of normally pathogen-induced salicylic acid signify the constitutively activated defense response against pathogen. This pattern is similar to that observed in other RNA processing factor mutants, such as *UPF* genes engaged in the nonsense-mediated decay (NMD) pathway, which also demonstrate constitutive resistance to *Pst*. The *upf* mutants also show upregulated expression of *PR* family genes, increased levels of salicylic acid, and reduced expression of *PDF1.2* (Jeong *et al*., 2011; Rayson *et al*., 2012).

We have also measured the levels of two other defense-related phytohormones, 12-oxo-phytodienoic acid (OPDA) and abscisic acid (ABA) (Fig. 1e). OPDA is a precursor of jasmonic acid but can also act as an independent signaling molecule (Wasternack & Strnad, 2016; Liu & Park, 2021; Jimenez Aleman *et al*., 2022), whereas ABA, acting antagonistically to SA and JA signaling, is mainly a negative disease resistance regulator but also controls callose-dependent responses and restricts pathogen entry (Melotto *et al*., 2008; Cao *et al*., 2011; Lim & Lee, 2015; Xin *et al*., 2018). Similarly to JA, a significant increase in OPDA accumulation was observed in DXO1-deficient plants (Fig. 1e). Since the metabolism of these two molecules is closely linked, it is possible that the final amount of JA in these plants is, at least to some extent, due to OPDA overproduction. In contrast, ABA levels in the *dxo1-2* line remained unchanged (Fig. 1e), consistent with no apparent effect on pathogen penetration of the mutant. Together, our results show that DXO1-deficient plants are characterized by an atypical biphasic susceptibility phenotype but ultimately exhibit constitutive immunity and increased resistance to *Pst* infection.

### DXO1 modulates PAMP-induced plant antibacterial defense

Pathogens that overcome the first line of defense are exposed to the plant’s innate immune system. Through the action of the PRR family receptors, plants detect pathogen-associated molecular patterns (PAMPs), which activate intracellular signaling pathways such as MAPK cascades, calcium influx, and ROS burst, triggering a wide range of defense responses in the plant (DeFalco & Zipfel, 2021; Ngou *et al*., 2022). Due to the observed initial susceptibility of the *dxo1-2* mutant to pathogen infection, we decided to examine the early PTI response of plants upon recognition of the conserved molecular patterns flg22 and elf18.

One of the earliest events elicited by PAMP perception involves the activation of at least two MAPK cascades: MEKK-MKK4/5-MPK3/6 and MEKK1-MKK1/2-MPK4, which play a positive or a negative role in pathogen defense, respectively (Petersen *et al*., 2000; Tena *et al*., 2011; Rasmussen *et al*., 2012; Meng & Zhang, 2013; Frei dit Frey *et al*., 2014; Lang *et al*., 2017). To assess the contribution of DXO1 to the activation of MAPKs upon PAMP treatment, we checked the levels of phosphorylated forms of MAPKs in the *dxo1-2* and wild-type seedlings incubated with flg22 and elf18. Immunoblotting with specific antibodies revealed that PAMP-triggered activation of all MAPKs was clearly delayed in the mutant, especially in the case of flg22, compared to wild-type plants (Fig. 2a). Thus, reduced activity of MAPKs in the absence of DXO1 may affect the functioning of many pathogen-related transcription factors via their delayed phosphorylation, particularly during the early stages of infection. This effect may account for the observed initial sensitivity of the mutant to pathogen infection.

**Fig. 2.**
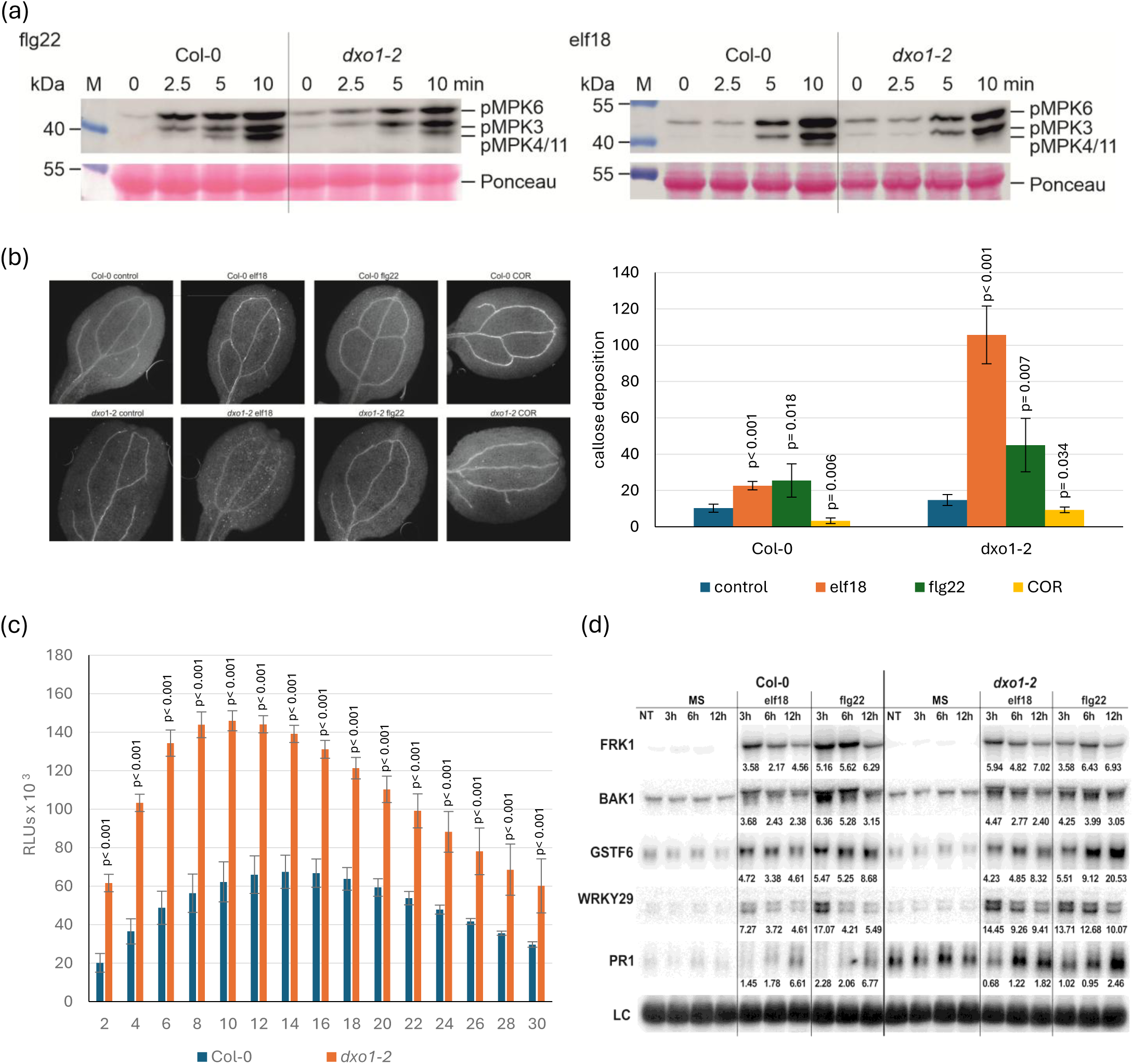
Pathogen-associated molecular patterns (PAMP)-induced activation of MAPKs, callose deposition, production of ROS, and expression of pathogenesis markers are altered in the *dxo1-2* mutant. (a) Detection of activated MPK3, MPK6 and MPK4/MPK11 kinases. Western blot analysis of protein extracts from 14-day-old Col-0 and *dxo1-2* plants at indicated time points after treatment with flg22 or elf18 (100 nM), using the anti-phospho-p42/44 MAPK antibody. M; protein size marker (molecular mass in kDa). Experiments were repeated at least three times; representative blots are shown. (b) One-week-old plants were treated with MS (control) or 1 mM of flg22, elf18 and COR. Callose formation was visualized by aniline blue staining and epifluorescence microscopy and quantified using ImageJ software from digital photographs as the number of local maxima specified by the average of RGB coloured pixels (callose intensity) in plant material. Bars represent the mean of three independent biological replicates with error bars showing SD; P < 0.05 (Tukey’s t-test). Representative pictures are shown. (c) ROS production in response to 100 nM flg22 treatment in leaf discs from 6-week-old Col-0 and *dxo1-2* plants. Bars represent the mean of four independent biological replicates with error bars showing SD, P < 0.05 for Tukey’s test. Luminescence is in Relative Light Units (RLUs). (d) Northern blot analysis of factors involved in the PAMPs response. Samples were collected at indicated time points from non-treated (NT) 14-day-old seedlings, treated with MS (control) or 100 nM of flg22 and elf18. The ratio of transcript level in treated Col-0 and *dxo1-2* relative to the control (MS) is shown. Values were normalized to 18S rRNA loading control (LC). Experiments were repeated three times; representative blots are shown.

Another early PTI response is the production of callose at the site of plant injury, which prevents pathogen penetration. In turn, coronatine (COR) phytotoxin produced by *Pst* promotes virulence by inhibiting callose deposition in the cell wall (Geng *et al*., 2014). To explore the effects of PAMPs on callose deposition, 7-day-old wild-type and mutant *dxo1-2* plants were treated with flg22, elf18, and COR effectors and examined by microscopy after 24 hours (Fig. 2b). We observed that callose deposition following flg22 and elf18 treatment was significantly increased in the *dxo1-2* mutant compared to Col-0. As expected, in the case of coronatine treatment, callose levels were lower than in control plants but still elevated in the *dxo1-2*. Notably, the expression of two genes encoding callose synthase, *GSL6* and *GSL7*, was significantly upregulated in the *dxo1-2* after *Pst* compared to Col-0 (Table S1). Increased callose deposition in the mutant suggests an enhanced response of the PTI defense system, which consequently limits the spread and multiplication of bacteria in the leaves.

PAMPs also contribute to the production of reactive oxygen species (ROS) as one of the first signaling molecules to trigger plant pathogen response pathways. To test whether DXO1 modulates activation of this early response, we measured ROS levels after flg22 treatment in the *dxo1-2* and wild-type plants using the luminol-based assay (Fig. 2c). A much stronger burst of apoplastic ROS observed in the *dxo1-2* line than in Col-0 may more effectively inhibit pathogen multiplication, contributing to the resistance of mutant to *Pst*.

Finally, we examined by northern blot the impact of the *dxo1* mutation on the PAMP-induced activation of selected PAMP-responsive mRNAs in 14-day-old seedlings treated with two major pathogen effectors, flg22 and elf18 (Fig. 2d). As for the *Pst* infection, an increased basal level of *PR1* mRNA was also observed in the *dxo1-2* untreated plants (Fig. 1b,c), confirming the constitutive resistance of the mutant. Although changes in activation of pathogenesis markers in the *dxo1-2* line triggered by PAMPs were less pronounced than in the case of *Pst* infection, the expression of *WRKY29* and *GSTF6* mRNAs at later time points for both flg22 and elf18 was significantly higher in the mutant than in Col-0 (Fig. 2d). This shows that, in contrast to the impact of DXO1 on the broad range of immune responses caused by the pathogen infection, it has a rather limited effect on the PAMP-induced response at the molecular level. In turn, DXO1 appears to more strongly modulate other steps of infection signaling, including ROS burst and activation of MAPK, which lead to later stages in the response to the pathogen.

### Expression of pathogenesis-related genes during *Pst* infection is deregulated in the absence of DXO1

Transcriptomic analysis of the *dxo1* mutant revealed a large number of differentially expressed genes, among which GO-terms related to biotic stresses were significantly enriched (Kwasnik *et al*., 2019; Pan *et al*., 2020). This prompted us to check how these pre-existing defects in gene expression profiles affect the ability to respond to infection. We performed 3′RNA-seq (Alpern *et al*., 2019) on 6-week-old Col-0 and the *dxo1-2* plants 48 hours after spraying with *Pst* DC3000 or MgCl_2_ as a control (Fig. 3, Database S1). The differential gene expression analysis revealed a significant number of affected genes between wild-type and mutant and between treatments. In the *dxo1*L*2* under control conditions (*dxo1-2*/MgCl_2_ vs Col-0/MgCl_2_), 2894 genes were upregulated, and 3900 genes were downregulated compared to Col-0, while after *Pst* (*dxo1*L*2*/*Pst* vs Col-0/*Pst*) 3129 genes were upregulated, and 4237 were downregulated (Fig. 3a,b). These results show that both the absence of DXO1 and *Pst* infection significantly impact gene expression, although to a different extent. 3′RNA-seq data also confirmed the changes in mRNA levels measured by northern blot (Fig. S3). The PCA plot suggests that the primary source of variability in this experiment is *dxo1* mutation (PC1) with a lesser effect of infection (PC2) (Fig. S4a). In agreement with this, *dxo1* mutation induces profound changes in gene expression observed in infected and mock-treated samples (Fig. S4b, Dataset S1, Table S1) (Kwasnik *et al*., 2019; Pan *et al*., 2020). Most importantly, a large number of genes with significantly altered expression in the *dxo1-2* plants belong to GO-terms associated with defense response. Specifically, of the 2654 upregulated and 2524 downregulated protein-coding genes 24.7% and 7.6%, respectively, represent defense response-related genes.

**Fig. 3.**
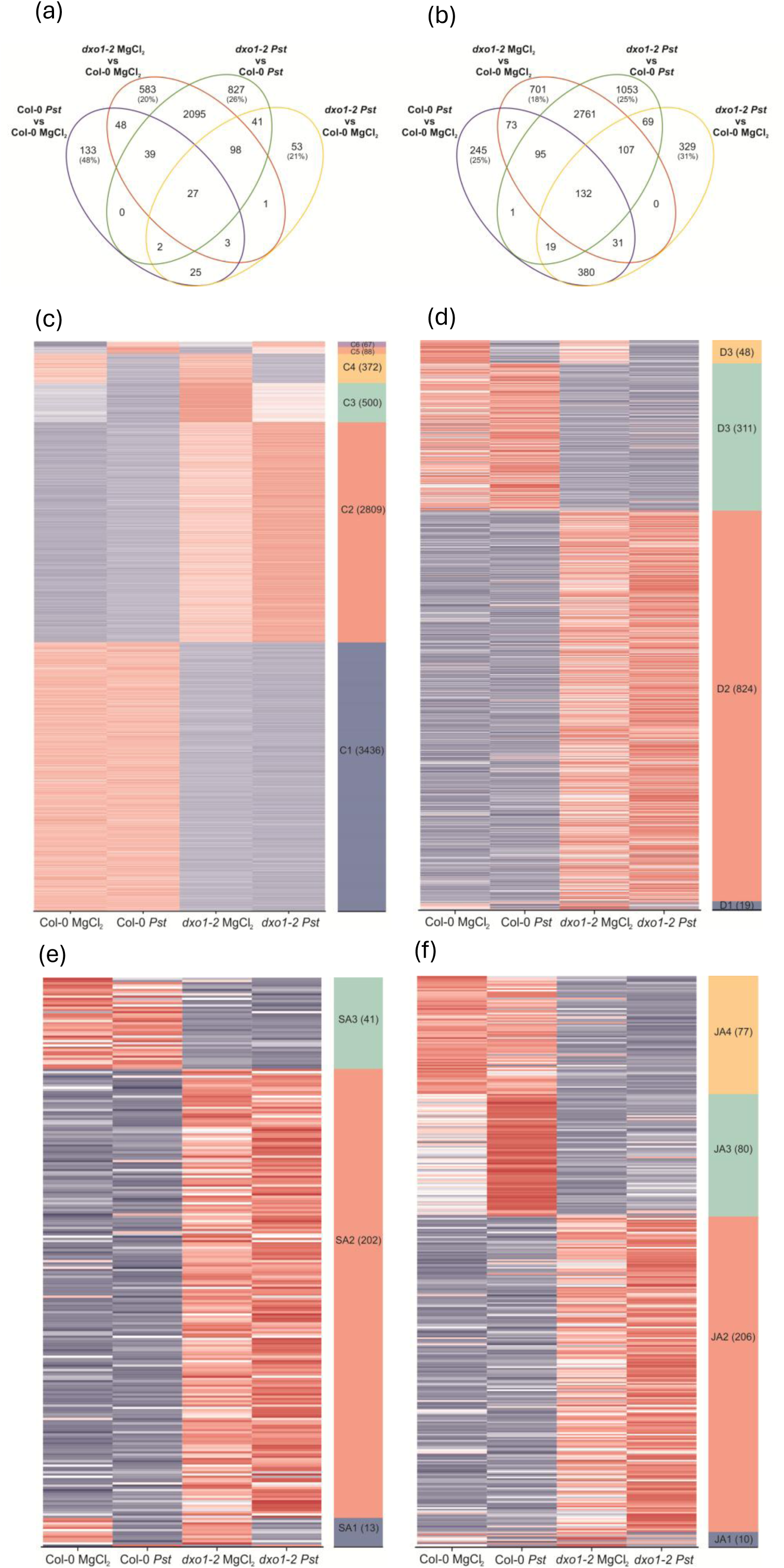
Genes affected by *Pst*-infection and lack of DXO1. (a, b) Venn diagrams indicate the number of significant overlapping upregulated (a) and downregulated (b) genes in Col-0 and the *dxo1-2* mutant in the control conditions and after *Pst* treatment. (c) Heatmaps illustrate the results of expression clustering for genes whose mRNA levels were significantly affected (DESeq2; absolute log2FC > 1; padj < 0.05) by *dxo1* mutation or the infection. (d-f) Heatmaps for gene expression clustering for genes selected as in (c) and narrowed down to specific functions in pathogen response. (d) Genes implicated in bacterial defense response (grouped in Gene Ontology terms: GO:0006952; GO:0042742; GO:0009617; GO:1900426; GO:0006968). (e) Genes implicated in SA response (grouped in Gene Ontology terms: GO:0009751; GO:0009697; GO:0009696; GO:0046244; GO:0009863; GO:2000031; GO:0080142; GO:0080151 GO terms). (f) Genes implicated in JA response (grouped in Gene Ontology terms: GO:0009753; GO:2000022; GO:0009694; GO:0009695). (c-f) Clustering was conducted using WGCNA and z-score normalised data. The number of genes included in each cluster is indicated on the right.

As expected from previous analyses, key pathogenesis-related genes *PR1*, *PR2*, *PR3*, *PR4*, and *PR5* were strongly upregulated in the *dxo1-2* mutant compared to Col-0 control, and even more activated after pathogen treatment (Table S1). Furthermore, consistent with the increased production of salicylic acid (see Fig. 1e), the expression of genes involved in the SA pathway, including major components such as *ICS1/SID2* (*ISOCHORISMATE SYNTHASE 1*/ *SALICYLIC ACID INDUCTION DEFICIENT 2*), *PAD4* (*PHYTOALEXIN DEFICIENT 4*), *CBP60G* (*CALMODULIN BINDING PROTEIN 60-LIKE G*), *PBS3* (*AvrPphB SUSCEPTIBLE 3*), and *EDS5* (*ENHANCED DISEASE SUSCEPTIBILITY 5*) and transcription factors *TCP8* (*TEOSINTE BRANCHED1/CYCLOIDEA/PCF*) and *TCP9* (Zhang & Li, 2019), was significantly elevated in the *dxo1-2* plants. In contrast, JA-regulated genes, e.g. *VSP1* (*VEGETATIVE STORAGE PROTEIN 1*), *VSP2*, *LOX2* (*LIPOXYGENASE 2*), and *PDF1.2* (Gupta *et al*., 2020) were highly downregulated. These observations further confirm that plants lacking DXO1 exhibit constitutive resistance to *Pst* and point to the involvement of DXO1 in the antagonistic interactions between SA and JA. This is corroborated by the increased level of major transcription factors that are involved in the JA-SA crosstalk, including *ANAC019*, *ANAC055*, *ANAC072* (*NAC DOMAIN CONTAINING PROTEIN*), *WRKY17*, *WRKY41*, *WRKY62*, and *WRKY70* (Pieterse *et al*., 2012; Yang *et al*., 2019).

SA, along with another defense hormone N-hydroxypipecolic acid (NHP), are two mobile signals leading to systemic and acquired resistance (SAR). Notably, in addition to SA pathway components, also genes involved in the NHP-dependent pathway, namely *ALD1* (*AGD2-LIKE DEFENSE RESPONSE PROTEIN 1*), *SARD4* and *FMO1* (*FLAVIN-DEPENDENT-MONOOXYGENASE 1*), as well as other SAR regulators (*SARD1* (*SAR DEFICIENT 1*), *NIMIN1* (*NIM1-INTERACTING 1*)) (Fu & Dong, 2013; Zeier, 2021), were upregulated in the mutant (Table S1). This raises the possibility that the *dxo1* mutation may lead to an enhanced SAR response.

Another interesting observation involves SERINE-RICH ENDOGENOUS PEPTIDES (SCOOPs), which belong to a group of plant hormones known as phytocytokines that regulate plant development and immunity (Del Corpo *et al*., 2024). Several SCOOP isoforms, derived from PROSCOOPs secreted peptide precursors, are unique to *Brassicaceae*, and many of them have been shown to be activated in the PTI response and regulate plant resistance to various pathogens, including *P. syringae* (Hou *et al*., 2021; Stahl *et al*., 2022; Yang *et al*., 2023; Wu *et al*., 2024; Zhang *et al*., 2024). In most described cases, SCOOP-triggered immunity is mediated via the SCOOP receptor kinase MIK2 (MALE DISCOVERER 1-INTERACTING RECEPTOR-LIKE KINASE 2) and coreceptors BAK1 and SERK4 (SOMATIC EMBRYOGENESIS RECEPTOR KINASE 4). SCOOP-induced activation of the MIK2-BAK1 complex has been shown to lead to a series of PTI responses, including activation of MAPK kinases and ROS burst (Hou *et al*., 2021; Rhodes *et al*., 2021). In the *dxo1-2* mutant, especially after *Pst* treatment, multiple SCOOP precursors (PROSCOOPs) were strongly upregulated (Table S1), most markedly for *PROSCOOP2*, followed by *PROSCOOP18, PROSCOOP17*, *PROSCOOP38*, *PROSCOOP19*, as well as the best characterized *PROSCOOP12*, which has been demonstrated to be involved in plant immunity (Gully *et al*., 2019; Zhai *et al*., 2024). Also, the expression of genes encoding other peptide phytocytokines, which contribute to the resistance to *Pst*, namely *IDL6 (INFLORESCENCE DEFICIENT IN ABSCISSION IDA-like 6)*, *GLV4/RGF7* (*ROOT MERISTEM GROWTH FACTOR 7*), *PIP2* (*PAMP-INDUCED SECRETED PEPTIDE 2*), *PSK4* (*PHYTOSULFOKINE 4*), and *RALF23* (*RAPID ALKALINIZATION FACTOR 23*) (Del Corpo *et al*., 2024), was altered in the *dxo1-2* mutant (Table S1). It is tempting to speculate that the resulting PTI response of DXO1-deficient plants may also be shaped by changes in defense pathways involving immunogenic phytocytokine peptides.

The strong effect of the *dxo1* mutation is also supported by the clustering of DEGs, in which pathogenesis-related genes affected exclusively in the mutant create the most prominent groups (clusters C1 and C2 in Fig. 3c). In turn, cluster C4 groups genes whose expression is similarly downregulated by infection, whereas clusters C3, C5, and C6 show differential responses to pathogen between the wild-type and the mutant. Of special interest is the large number of genes in GO terms corresponding to defense and the defense-related hormones JA and SA (clusters 2 and 3 in Fig. 3d-f). In particular, cluster 2 contains genes upregulated in the *dxo1-2* mutant, which have even more enhanced expression upon infection. This concerns for example *PR1*, *PR2*, *PR5*, *ANAC019*, *ANAC055*, *ANAC075*, *NUDIX HYDROLASE HOMOLOG* (*NUDT4*, *NUDT5*, *NUDT6*, *NUDT17*, *NUDT18*, *NUDT21*, *NUDX25*), and *WRKY* transcription factors (*WRKY17*, *WRKY18*, *WRKY38*, *WRKY41*, *WRKY46*, *WRKY53*, *WRKY60*, *WRKY67*, *WRKY*70), as well as *NIMIN1*, *NIMIN2*, *FMO1*, *GRX480*, *JAZ8*, *JAZ11*, *LOX1*, *MYB2* (*MYB DOMAIN PROTEIN 2*), *MPK4* (*MAP KINASE 4*), *RPS2* (*RESISTANT TO P. SYRINGAE 2*), *EDS5* (Dataset 2, Table S1). Moreover, clustering of the defense-, SA- and JA-related gene groups to obtain a more detailed picture focused solely on changes in the expression of genes associated with response to infection revealed a similar trend, i.e. increased expression in the *dxo1-2* mutant, further enhanced after biotic stress (Fig. 3d-f).

Finally, to assess the contribution of the catalytic site and the N-terminal domain of DXO1 to changes in expression of genes belonging to GO terms related to biotic stress response, we reanalysed the 3′RNA-seq data for the four transgenic lines *dxo1-2*+DXO1(WT), *dxo1-2*+DXO1(E394A/D396A), *dxo1-2*+DXO1(ΔN194), and *dxo1-2*+DXO1(ΔN194/E394A/D396A) (Zakrzewska-Placzek *et al*., 2025) and compared the results with those for Col-0 and the *dxo1-2* under control conditions (Dataset 2, Fig. 4). In the defense response GO term, as expected, the expression pattern in Col-0 and dxo1-2+DXO1(WT) is similar for all clusters, except D3 and D7 (Fig. 4a). Moreover, a significant number of genes in different clusters (D2, D4, D5) have comparable expression in the *dxo1-2* as in the two lines expressing DXO1 variant lacking the N-terminal domain, *dxo1-2*+DXO1(ΔN194) and *dxo1-2*+DXO1(ΔN194/E394A/D396A), but not in *dxo1-2*+DXO1(E394A/D396A). The largest group with similarly increased expression occurs in D2 and contains key pathogenesis-related genes *PR1*, *PR2*, *PR4*, *PR5*, *WRKY* transcription factors (*WRKY18*, *WRKY33*, *WRKY40*, *WRKY54*, *WRKY60*, *WRKY*70) and *NIMIN1*, *NIMIN2* and *NIMIN3*. In contrast, D4 and D5 consisting of genes with comparably reduced expression, include, for example, factors involved in the JA pathway, such as *JAZ9*, *JAZ3*, *JAZ9*, *JAZ10*, *JAZ13*, and *PDF1.2A*, *PDF1.2B*, *PDF1.2C*, *PDF1.3*, *PDF2.1*, and *PDF2.2*. Thus, the expression of up to 66% of genes in the defense GO terms shows dependence on the DXO1 N-terminal domain and not on the catalytic site. Also, genes in clusters JA2, JA3, JA5 and SA2, SA4, SA5, in JA- and SA-related GO terms, respectively, have a similar expression pattern in mutants expressing DXO1 lacking the N-terminal domain (Fig. 4b,c). However, there are some less abundant clusters in the SA pathway GO terms that rely solely on the catalytic activity of DXO1 (cluster SA3) or on both the catalytic activity and the N-terminal domain (cluster SA4). This gene cluster analysis reinforces the notion that most of the changes in the expression of factors regulating the antibacterial defense response induced by the *dxo1* mutation are primarily associated with the N-terminal domain of the protein. It should be noted, however, that some aspects of this response also depend on the enzymatic activity of DXO1, highlighting the complex function of this protein in plant immunity.

**Fig. 4.**
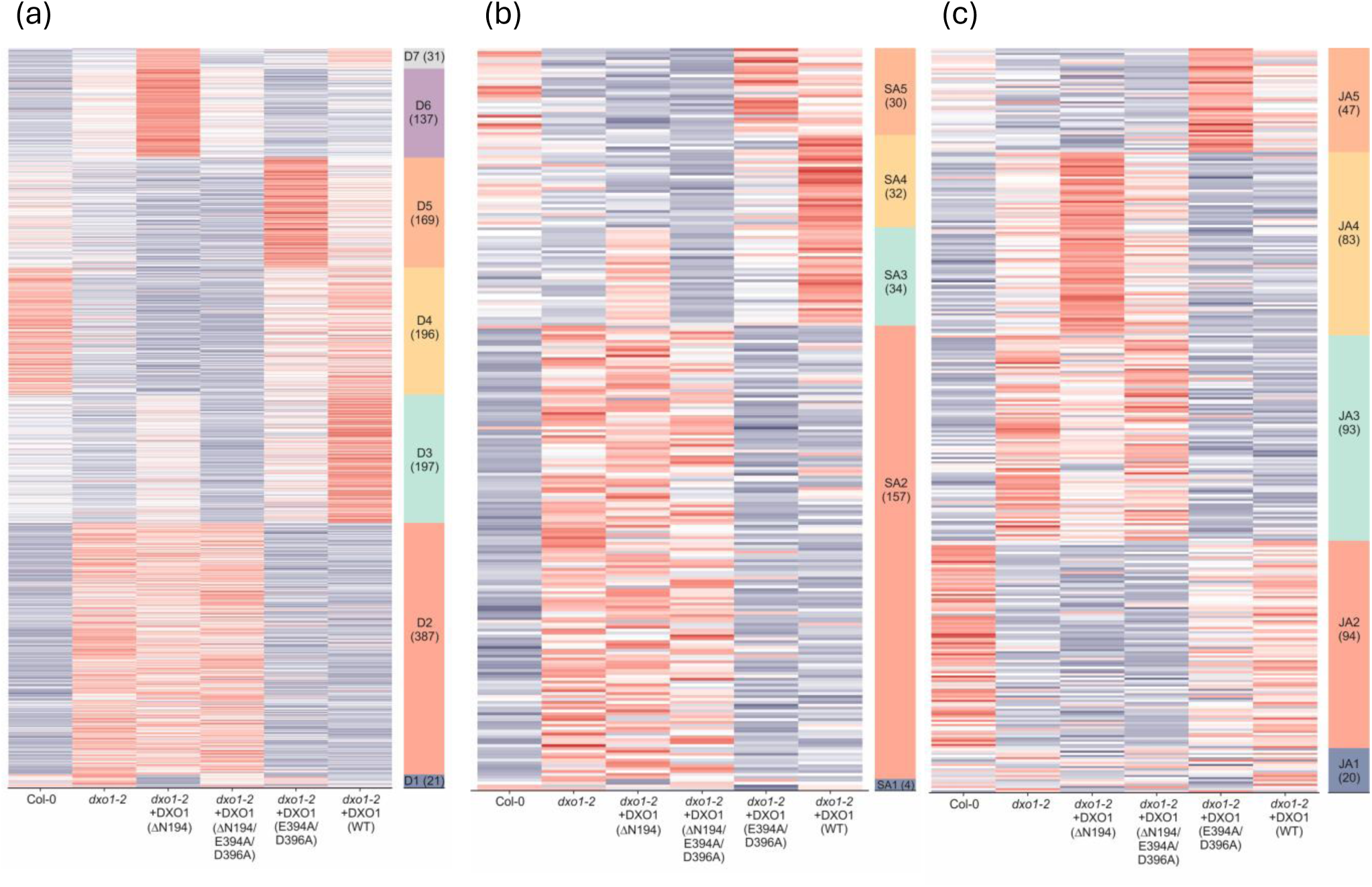
Heatmaps illustrate the results of expression clustering for genes whose mRNA levels were significantly affected (DESeq2; absolute log2FC > 1; padj < 0.05) in plant lines with a mutated DXO1 gene, complemented by genetic constructs containing different versions of DXO1 (GSE210631; Zakrzewska et al., 2025). Affected genes were narrowed down to specific functions in pathogen response: (a) Genes implicated in bacterial defence response (grouped in Gene Ontology terms: GO:0006952; GO:0042742; GO:0009617; GO:1900426; GO:0006968); (b) Genes implicated in SA response (grouped in GO terms: GO:0009751; GO:0009697; GO:0009696; GO:0046244; GO:0009863; GO:2000031; GO:0080142; GO:0080151); (c) Genes implicated in JA response (grouped in Gene Ontology terms: GO:0009753; GO:2000022; GO:0009694; GO:0009695). (a-c) Clustering was conducted using WGCNA and z-score normalised data, with the number of genes included in each cluster indicated on the right.

### Both the catalytic site and the plant-specific N-terminal extension of DXO1 contribute to the response to *Pst* infection

Most of the molecular and morphological defects observed so far in the *dxo1* mutants depended on the plant-specific N-terminal domain (Kwasnik *et al*., 2019; Carpentier *et al*., 2025). To check whether this also applies to the pathogen response and to independently assess the contribution of the catalytic site and the N-terminal extension (NTE), we used transgenic *dxo1-2* lines expressing different DXO1 variants: DXO1(WT) control, DXO1(E394A/D396A) catalytic mutant and DXO1(ΔN194) lacking the NTE (Kwasnik *et al*., 2019). After *Pst* treatment, all transgenic lines showed sensitivity to the pathogen at 24 hpi compared to WT and were resistant at 48 and 72 hpi (Fig. 5a,b). The response of the transgenic mutants closely resembles the behavior of the *dxo1-2* plants exhibiting initial sensitivity followed by resistance, confirming that this is a genuine phenotype associated with DXO1 deficiency. More importantly, these results indicate that resistance to the pathogen depends not only on the N-terminal extension, as for other *dxo1*-related phenotypes described previously, but also on the active catalytic domain. This is the first observation showing the involvement of the DXO1 enzymatic activity in plant physiology, specifically in the response to biotic stress.

**Fig. 5.**
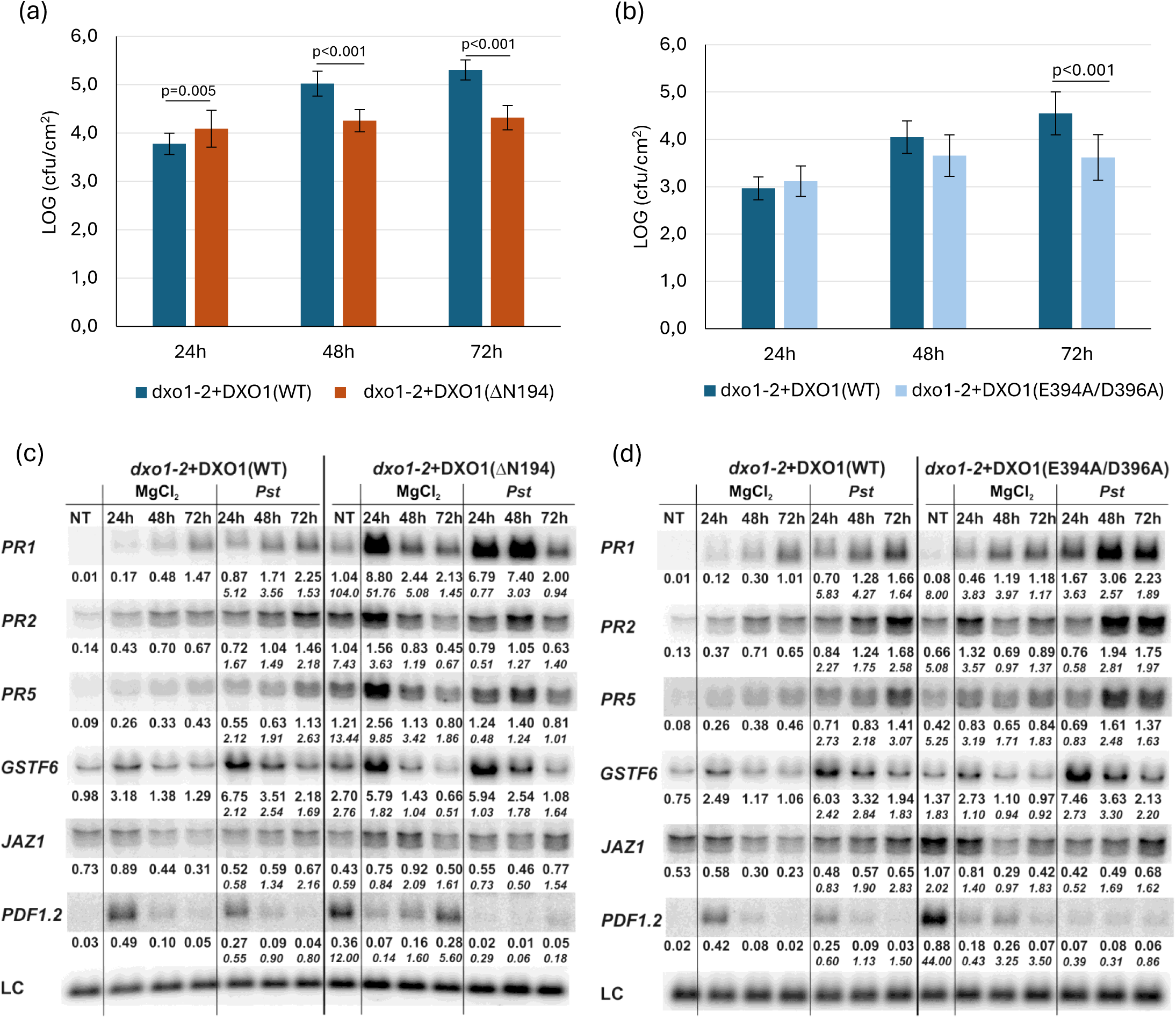
The transgenic lines are resistant to *Pseudomonas syringae* pv. *tomato* DC3000. (a, b) Growth of *Pst* DC3000 after 24, 48, and 72 h in *dxo1-2*+DXO1(WT) and *dxo1-2*+DXO1(ΔN194) (a) or *dxo1-2*+DXO1(WT) and *dxo1-2*+DXO1(E394A/D396A) (b). For each time point, leaf discs were collected from 6 plants. Results are the mean of four independent experiments, and error bars represent SD; P < 0.05 (Tukey’s t-test). (c, d) Northern blot analysis of factors involved in response to *Pst* DC3000. Samples were collected from non-treated (NT), control (MgCl_2_) and infected (*Pst*) *dxo1-2*+DXO1(WT) and *dxo1-2*+DXO1(ΔN194) (c) or *dxo1-2*+DXO1(WT) and *dxo1-2*+DXO1(E394A/D396A) (d). Numbers represent the ratio of transcript level in treated transgenic lines versus *dxo1-2*+DXO1(WT) normalized to U2 snRNA loading control (LC), which is shown as the main numbers, while the ratio relative to the control conditions is given in italics.

Analysis of the pathogen response at the molecular level for transgenic lines, based on the examination of the main pathogenesis markers by northern blot, showed enhanced expression of *PR* genes and downregulation of *PDF1.2* in both mutants compared to the WT control (Fig. 5c,d), similarly as observed for the *dxo1-2* (see Fig. 1c). This further confirmed the involvement of both DXO1 features, NTE and catalytic activity, in this process. However, there were also significant differences between the two transgenic lines expressing DXO1 variants. The DXO1(ΔN194) mutant showed strong activation of *PR* genes, especially *PR1* and *PR5*, already in untreated plants and control conditions, as well as at an early stage of infection (i.e. 24 hpi) as compared to DXO1(WT) and DXO1(E394A/D396A), for which activation of these genes was observed at 48 hpi and 72 hpi. This pattern is reminiscent of the effect seen in the *dxo1-2* plants and reflects the constitutive defense response for both the *dxo1-2* and DXO1(ΔN194) lines. Notably, the autoimmunity of the *dxo1* plants appears more pronounced when the DXO1 variant lacking the NTE is expressed than in the *dxo1* knockout, raising the possibility that the truncated protein acts in a dominant-negative manner. In contrast, expression of *PR* genes in the DXO1(E394A/D396A) line was stronger than in DXO1(WT) but followed the same pattern and reached the highest level also at 48 and 72 hpi.

Consistently, SA, JA and OPDA were strongly increased only in DXO1(ΔN194) plants, as was the case for the *dxo1-2* mutant, but not in DXO1(E394A/D396A) (Fig. 6a). At the same time, ROS production following flg22 treatment was higher in both transgenic mutants (Fig. 6b). Together, these observations support the involvement of both the N-terminal domain of DXO1 and catalytic activity in the response to biotic stress and suggest that the catalytic center is not directly involved for the autoimmunity phenotype of *dxo1* mutants.

**Fig. 6.**
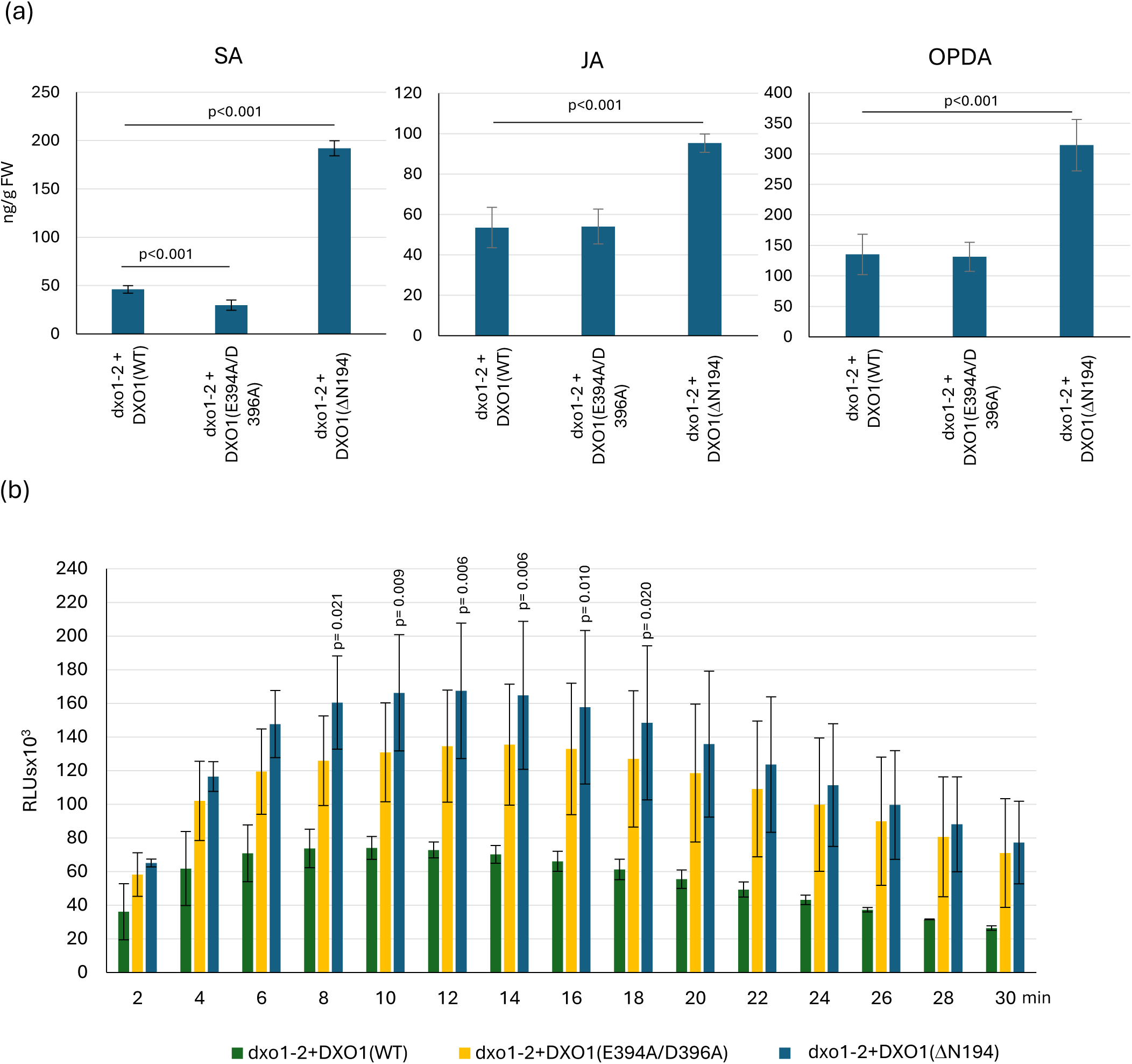
The transgenic lines are involved in stress response. (a) The concentration of SA, JA, and OPDA in 14-day-old seedlings of *dxo1-2*+DXO1(WT), *dxo1-2*+DXO1(E394A/D396A), and *dxo1-2*+DXO1(ΔN194). Bars represent the mean of three independent biological replicates with error bars showing SD, P < 0.05 for Tukey’s test. (b) ROS production in response to 100 nM flg22 treatment in leaf discs from 6-week-old *dxo1-2*+DXO1(WT), *dxo1-2*+DXO1(E394A/D396A), and *dxo1-2*+DXO1(ΔN194) plants. Bars represent the mean of three independent biological replicates with error bars showing SD, P < 0.05 for Tukey’s test. Luminescence is in Relative Light Units (RLUs).

### DXO1 influences the response to biotic stress by regulating the stability of pathogenesis-related mRNAs

The specific contribution of DXO1 enzymatic activity to the regulation of plant resistance to pathogen infection prompted us to investigate the involvement of DXO1 in the degradation of mRNAs encoding pathogenesis-related factors. It was recently reported that catalytic inactivation of DXO1 impairs CTRD (co-translational mRNA decay), which is an XRN4- dependent mechanism of 5’-3’ degradation of mRNAs still undergoing translation on polysomes (Merret *et al*., 2013, 2015; Carpentier *et al*., 2020, 2025). Moreover, we previously observed stabilization of a number of mRNAs in *xrn4-5* and *dxo1-2* mutants, including those involved in the response to biotic stress (Golisz *et al*., 2013; Kwasnik *et al*., 2019). Therefore, we investigated the half-life of several pathogen-related mRNAs and we observed that they were also stabilized in the absence of DXO1 (Fig. 7a). Finally, we established that the *xrn4-5* mutation results in increased resistance to *Pst* infection after 72hpi and alters the expression of selected major pathogenesis markers (Fig. 7b,c). These findings suggest that the enzymatic role of DXO1 and XRN4 in pathogen response may be linked to degradation of mRNAs encoding defence-related factors, via general mRNA decay or the CTRD mechanism.

**Fig. 7.**
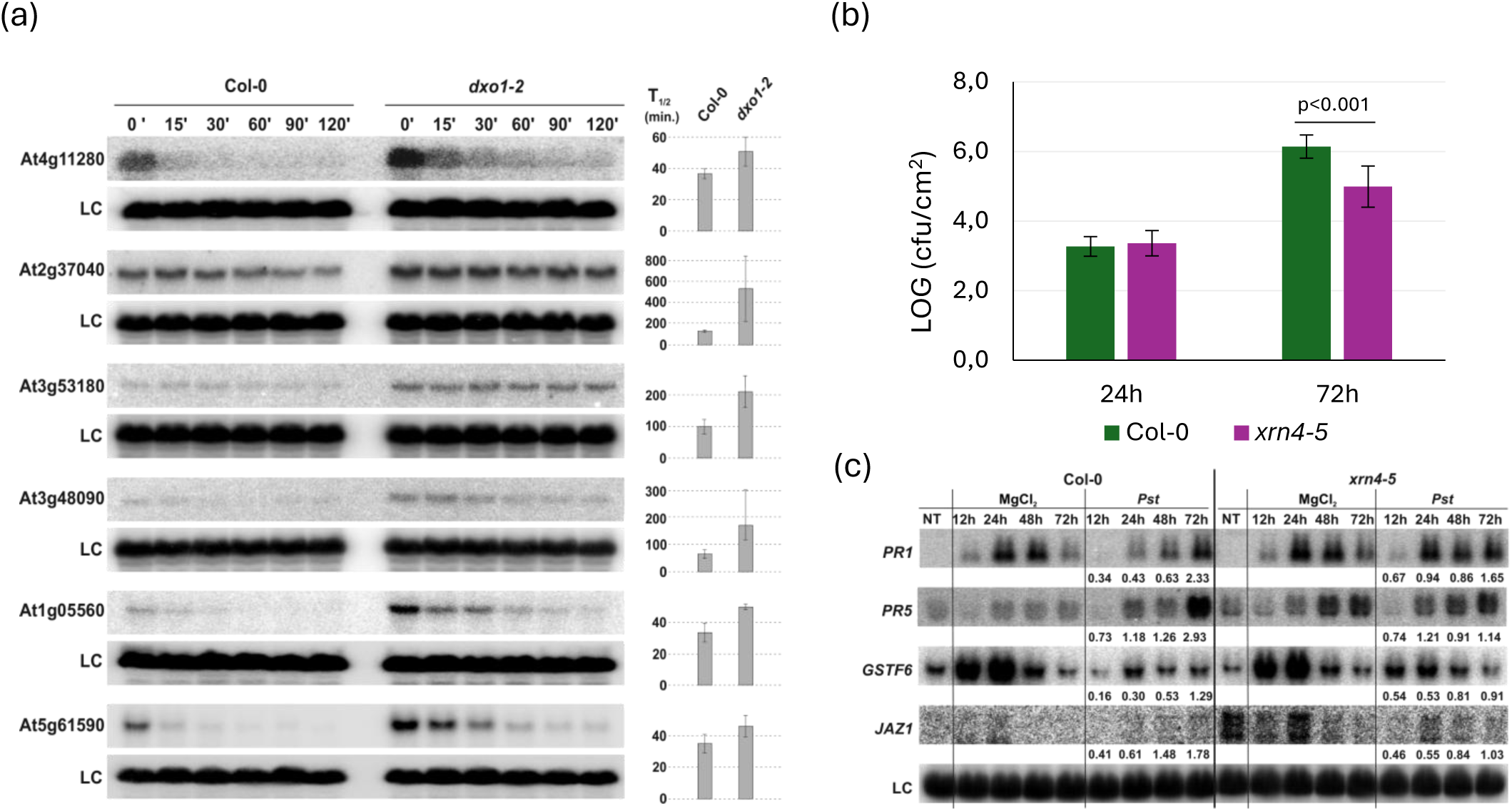
DXO1 affects mRNA stability. (a) Northern blot analysis of mRNAs involved in pathogen response at indicated time points after cordycepin treatment in Col-0 and *dxo1-2* plants. The estimated mRNA half-life (T_1/2_) was calculated by regression of the semi-logarithmic mRNA levels against time. Mean T_1/2_ values calculated from three biological replicates are shown on the right of each panel with error bars representing SD. 18S rRNA was used as a loading control (LC). The *xrn4-5* mutant is resistant to *Pseudomonas syringae* pv. *tomato* DC3000. (b) Growth of *Pst* after 24 and 72 h in Col-0 and the *xrn4-5* mutant after spraying. For each time point, leaf discs were collected from 6 plants. Results are the mean of three independent experiments, and error bars represent SD; P < 0.05 (Tukey’s t-test). (c) Northern blot analysis of factors involved in response to *Pst* DC3000. Samples were collected from non-treated (NT), control (MgCl_2_) and infected (*Pst*) Col-0 and the *xrn4-5* plants at the indicated time points. Numbers represent the ratio of transcript level in treated *xrn4-5* versus Col-0 normalized to 18S rRNA loading control (LC). Experiments were repeated at least three times; representative blots are shown.

## Discussion

Arabidopsis DXO1 is the sole plant homolog of the DXO protein family. Its structural features are markedly different from those of fungal and mammalian homologs, resulting in distinct activities and cellular functions across various organisms. This structural and functional distinction reflects specialized evolutionary adaptation that endows Arabidopsis DXO1 with unique roles in plant biology, including involvement in chloroplast metabolism and stress response pathways (Kwasnik *et al*., 2019; Pan *et al*., 2020; Yu *et al*., 2021; Zakrzewska-Placzek *et al*., 2025; this work). In this work, we confirmed the involvement of Arabidopsis DXO1 in plant immunity by showing that DXO1-deficient plants are resistant to *P. syringae* infection, but we also observed that this resistance develops in the later phase, after an initial hypersensitivity. We also showed that the constitutive defense response phenotype of the *dxo1* mutant is associated with the activation of the salicylic acid pathway and defense-related genes, which, together with the upregulation of genes involved in the N- hydroxypipecolic acid (NHP) signaling, probably leads to enhanced systemic and acquired resistance (SAR) response of the mutant plants. Finally, our data support the key contribution of the plant-specific N-terminal domain of DXO1 to the regulation of plant immunity, but also indicate the involvement of the enzymatic activity of the protein in this process.

An intriguing observation is that *dxo1-2* plants are initially sensitive to *Pst* but acquire final resistance only at a later stage, which is consistent with earlier studies showing enhanced resistance of the mutant three days after infection (Pan *et al*., 2020). The primary susceptibility of *dxo1-2* following surface inoculation, as opposed to direct injection, indicates that the more invasive method leads to the development of a faster and more robust defence. This effect can be attributed to the distinct morphology of *dxo1-2* leaves, namely their thinner epidermis, which normally acts as a first barrier against pathogen invasion (Javelle *et al*., 2011; Underwood, 2012), and the increased density of trichome, which can be used by bacterial pathogens as colonization or entry sites, especially through damaged parts such as broken trichome bases (Mansvelt & Hattingh, 1987; Beattie & Lindow, 1999; Kim, 2019). In addition, the initial sensitivity of the mutant is also manifested by delayed phosphorylation of MAP kinases following PAMP treatment. This may, in turn, cause a delay in the activation of different transcription factors, resulting in a slower response to infection. These observations highlight the complexity of plant-pathogen interactions and point to the still underappreciated role of leaf morphology in determining susceptibility and resistance.

The most striking hallmark of plant DXO1 is its unique, large N-terminal extension (NTE) that stabilizes protein-RNA interactions (Kwasnik *et al*., 2019; Pan *et al*., 2020). Through this domain, DXO1 interacts with and activates RNA guanosine-7 methyltransferase RNMT1, which ensures participation of DXO1 in mRNA cap biogenesis (Xiao *et al*., 2023). In addition, the NTE is essential for the chloroplast-related functions of DXO1, even though this protein is not found in chloroplasts. Also other molecular and morphological defects in *dxo1-2* mutants, including the altered expression of genes involved in photosynthesis, accumulation of RNA quality control siRNAs (rqc-siRNAs), chloroplast functioning, defense response, and ribosome maturation and function, mainly depend on the plant-specific extension, as these phenotypes are complemented by the expression of the catalytically inactive DXO1(E394A/D396A) variant, but not by the truncated DXO1(ΔN194) protein (Kwasnik *et al*., 2019; Pan *et al*., 2020). Since NTE is responsible for the DXO1-RNMT1 interaction, one possible explanation for its key contribution to cellular activities of DXO1 could be the fine-tuning of mRNA expression levels by regulating their proper capping, as is probably the case for ribosome biogenesis genes (Kwasnik *et al*., 2019; Pan *et al*., 2020). However, this function of NTE cannot fully account for its role in pathogen response, because *dxo1-2* and *rnmt1-2* mutations have rather opposite effects on defense-related genes that are generally upregulated in *dxo1-2* plants and are unaffected or downregulated in the *rnmt1-2* mutant (Kwasnik *et al*., 2019; Pan *et al*., 2020). Therefore, another action of NTE, for example mediating interactions with plant immunity regulators, may additionally determine its significant input into this process.

Our research provides evidence for an enzymatic role of DXO1 in physiological processes in Arabidopsis. We have shown for the first time that the plant response to *Pst* depends not only on the N-terminal extension but also on the active catalytic domain of DXO1. The transgenic *dxo1-2* lines expressing various DXO1 variants, including the catalytically inactive DXO1(E394A/D396A), and the truncated DXO1(ΔN194), exhibited similar susceptibility to *Pst* infection as the *dxo1-2* mutant. This indicates the essential role of both structural features of DXO1 in regulating the immune response. Additionally, the clustering of defense-, SA- and JA-related gene groups, differentially expressed in transgenic lines, not only confirmed the important contribution of the DXO1 NTE but also revealed a number of clusters in SA pathway that depend on the catalytic domain. One of our most important findings is that the two unique properties of the DXO1 protein, the N-terminal extension and the catalytic activity, represent distinct aspects of the pathogen response pathway. It appears that the lack of the NTE, rather than the catalytic residues, is responsible for the SA-dependent autoimmunity phenotype of *dxo1* mutant plants. All these observations clearly underscore the critical role of the N-terminal extension and raise intriguing questions regarding the significance of DXO1 enzymatic activity in the context of plant biology.

DXO1 has been shown *in vitro* to be primarily a deNADding enzyme, efficiently removing NAD cap from NAD-capped RNAs but unable to hydrolyze the m^7^G or the unmethylated cap, and additionally exhibiting a weak 5′ to 3′ exoribonuclease activity (Kwasnik *et al*., 2019; Pan *et al*., 2020). It is possible, however, that in plant cells DXO1 has a wider range of possible functions through interactions with other proteins. A recent report demonstrated that catalytic activity of DXO1 is required for co-translational mRNA decay (CTRD) (Carpentier *et al*., 2025), a XRN4-dependent 5’-3’ degradation mechanism of mRNAs that are still being translated on polysomes (Merret *et al*., 2013, 2015; Carpentier *et al*., 2020, 2025). Considering the deNADding activity of DXO1, the activation of CTRD under stress conditions, and the enrichment of DXO1 CTRD targets in GO terms linked to stress response, it was proposed that DXO1 plays a specific role in this pathway by targeting NAD^+^-capped mRNAs, particularly those related to abiotic stress (Carpentier *et al*., 2025). However, it cannot be ruled out that CTRD is also involved in modulating the response to other types of stress, including biotic stress. We have shown that a number of pathogen response factor mRNAs are stabilized in the *dxo1-2* mutant (Kwasnik *et al*., 2019; this work). Importantly, mRNAs involved in the biotic stress response are also stabilized in the *xrn4-5* mutant (Golisz *et al*., 2013), which is resistant to *Pst* (this work). It can therefore be assumed that the enzymatic activity of DXO1, and perhaps also XRN4, contributes to the regulation of plant immunity through their role in the CTRD mechanism.

Plant stress responses, including disease resistance, are closely linked to chloroplast function. Chloroplasts produce a variety of signals involved in antibacterial defense such as ROS, Ca ^2+^ ions, phytohormones, and metabolites that regulate retrograde signaling (e.g. PAP) (Serrano *et al*., 2016; Sowden *et al*., 2017; Kachroo *et al*., 2021). The impact of the DXO1 protein on the proper functioning of these organelles should also be considered. Although DXO1 is absent from chloroplasts, it is essential for chloroplast-related functions. The *dxo1-2* mutants exhibit severe growth retardation, low fertility, pale green coloration, and reduced chlorophyll content (Kwasnik *et al*., 2019; Pan *et al*., 2020; Xiao *et al*., 2023). Moreover, this mutation results in the downregulation of genes involved in photosynthesis and chloroplast functioning (Kwasnik *et al*., 2019; Pan *et al*., 2020; Xiao *et al*., 2023). Finally, DXO1 knockout affects pre-rRNA processing and ribosome biogenesis not only in the nucleolus but also in chloroplasts (Zakrzewska-Placzek *et al*., 2025). DXO1 is associated with chloroplasts via the 3′- phosphoadenosine-5′-phosphate (PAP) signaling pathway (Kwasnik *et al*., 2019). PAP, a chloroplast retrograde signal, accumulates under high light and drought stress and modulates RNA metabolism by inhibiting the activities of XRN 5′-3′ exoribonucleases and DXO1 (Estavillo *et al*., 2011; Kwasnik *et al*., 2019). In addition, DXO1 and XRN enzymes are controlled by a shared mechanism involving PAP regulators. One of these regulators is FRY1, a nuclear-encoded protein localized in chloroplasts and mitochondria, which dephosphorylates PAP (Estavillo *et al*., 2011). FRY1 is also involved in plant immunity by regulating SA- and JA-mediated signaling pathways (Ishiga *et al*., 2017). Finally, FRY1 has recently been shown to influence CTRD by inhibiting XRN4 and DXO1, most likely via PAP accumulation (Carpentier *et al*., 2025).

Integrating our findings with previous studies, we conclude that DXO1 regulates the response to biotic stress at the transcriptional level to fine-tune the expression of pathogenesis-related genes, as well as at the post-transcriptional level, possibly via the CTRD pathway, to modulate their stability and perhaps also translation (Fig. 8).

**Fig. 8.**
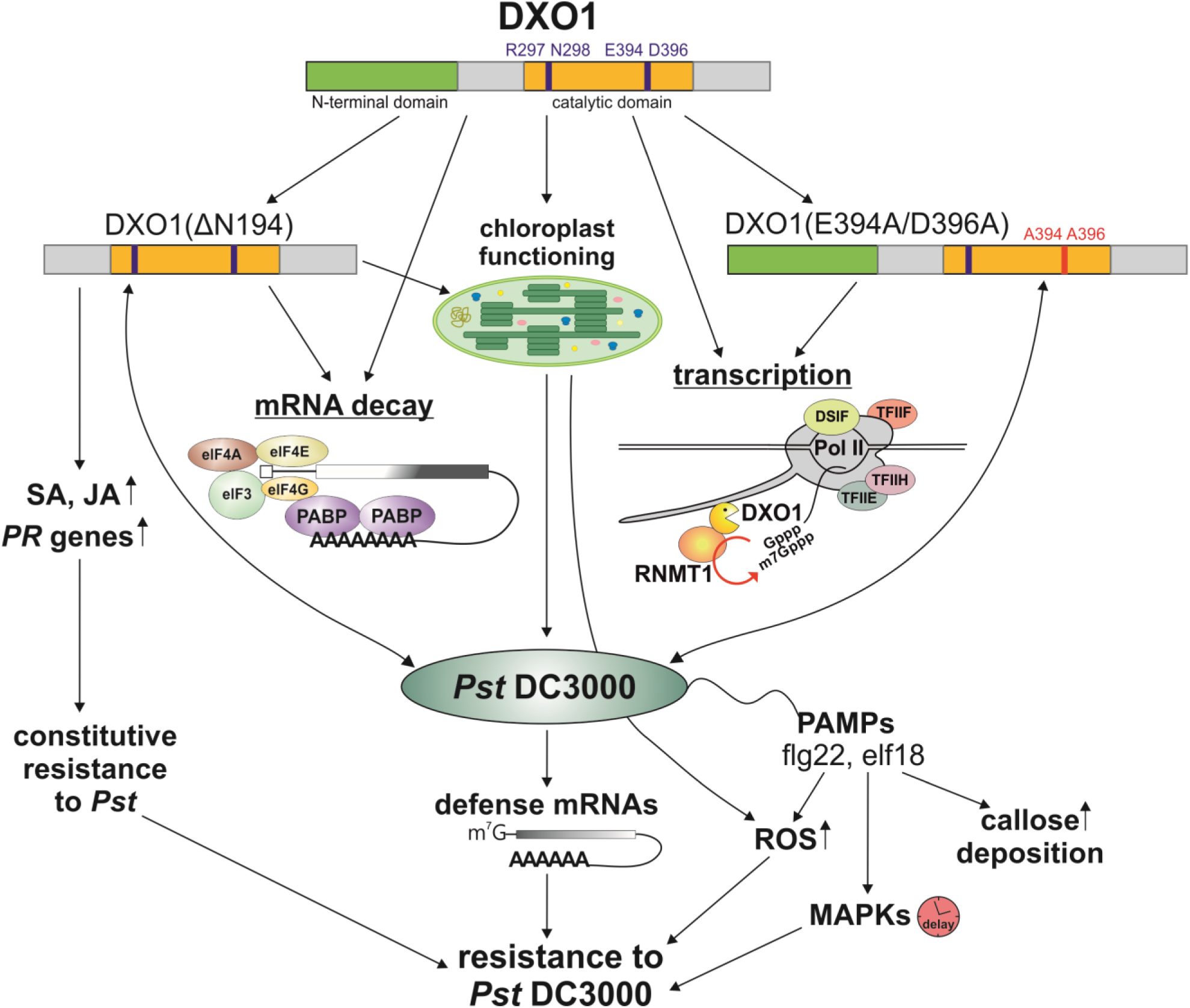
DXO1 affects several aspects of plant immunity. DXO1-deficient plants are resistant to *Pst* infection and show marked changes in the expression of key pathogenesis markers. This response depends not only on the N-terminal extension but also on the active catalytic site of DXO1. Plants expressing the full-length DXO1 and the variant lacking the N-terminal domain, but not the catalytic mutant, exhibit constitutive resistance to *Pst* due to increased levels of *PR* genes, SA and JA. Following PAMP treatment other markers of plant immunity, such as callose deposition and production of reactive oxygen species, are strongly induced, but the phosphorylation of MAP kinases is delayed. DXO1 is also relevant for disease resistance through chloroplast-related functions.

## Acknowledgements

The equipment used was sponsored in part by the Centre for Preclinical Research and Technology (CePT), a project co-sponsored by the European Regional Development Fund and Innovative Economy, The National Cohesion Strategy of Poland. This work was supported through grants from National Science Centre UMO-2014/13/B/NZ3/00405 to AG-M, UMO- 2018/29/B/NZ3/01980 and UMO-2021/40/Q/NZ1/00014 to JK.

## Author contributions

AG-M designed and performed most of the experiments and conceived the project and wrote the manuscript with MZ-P and MK contributions. MK analyzed the 3′RNA-seq results. MZ-P, ND, JP and WK performed some of the experiments. AM-C performed microscope analysis. JK supervised and completed the writing. All authors contributed to the article and approved the submitted version.

## Data availability

The data that support the findings of this study are publicly available. 3′RNA-seq data are deposited in the Gene Expression Omnibus database under accession codes GSE210631 (different complementation lines of *dxo1-2* mutant) and GSE291552 (Col-0 and *dxo1-2* upon control MgCl_2_ or *Pst* treatment).

## Supporting Information

**Dataset S1** List of significantly affected genes based on 3′ RNA-seq.

**Dataset S2** List of genes in clusters in Col-0 and *dxo1-2*.

**Dataset S3** List of genes in clusters for different complementation lines of *dxo1-2* mutant.

**Fig. S1** Cross-section of Arabidopsis leaves. (a, c) Col-0, (b, d) *dxo1-2*, (a, b) leaves at the vein, (c, d) leaves structure. Red lines show the border between palisade and spongy mesophyll cells. AdE— adaxial epidermis, AbE—abaxial epidermis, PM—palisade mesophyll, SM—spongy mesophyll, VT— vascular tissue. Bars represent the mean of three independent biological replicates with error bars showing SD, P < 0.05 for Tukey’s test.

**Fig. S2** Leaf morphology in Col-0 and *dxo1-2* plants. (a) Stomata density and aperture length on the leaf surface of 6-week-old Col-0 and *dxo1-2* plants were estimated using a Nikon Eclipse 80i microscope (Nikon). Scale bar, 25 µm. Bars represent the mean of five independent biological replicates with error bars showing SD, P < 0.05 for Tukey’s test. (b) Trichome density on the leaf surface of 6-week-old Col-0 and the *dxo1-2* plants. Images of the abaxial and adaxial leaf surfaces were captured using a digital microscope (VHX-7000N Keyence). Each trichome is circled in red (adaxial) or yellow (abaxial) to facilitate comparison of trichome density. Four leaves were analyzed for each genotype. Bars represent the mean of three independent biological replicates with error bars showing SD, P < 0.05 for Tukey’s test.

**Fig. S3** Comparison of the expression level of selected pathogen response genes in Col-0 and the *dxo1-2* mutant based on RNA-seq and northern blot analysis. Numbers represent fold change.

**Fig. S4** Results of RNA-seq analysis. (a) PCA (Principal component analysis) shows that biological replicas in 3′RNA-seq create four groups based on the presence of the *dxo1-2* mutation and *Pst* infection. (b) Volcano plots depicting the fold change in mRNA levels with comparisons between different conditions indicated above each plot. Significantly affected mRNAs (DESeq2; padj < 0.05) are represented by colours: upregulated (log2FC > 1 – red) and downregulated (log2FC < -1 – blue).

**Table S1** List of genes involved in different aspects of pathogen response in the *dxo1-2* mutant.

**Table S2** List of primers used in this study.

