## Supplemental Figures for "The Arabidopsis deNADding enzyme DXO1 modulates the plant immunity response"

**Fig. S1**

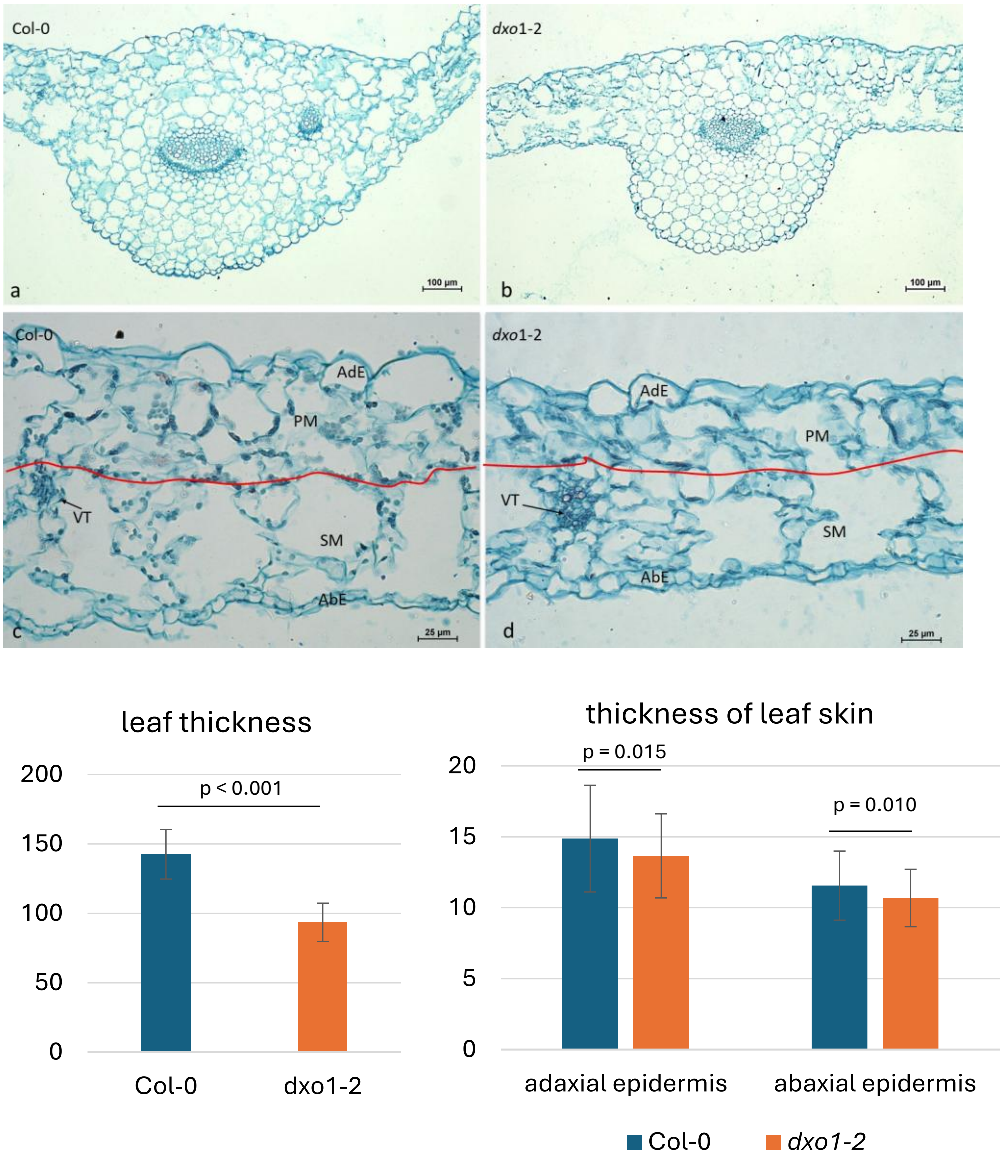


**Fig. S1** Cross-section of Arabidopsis leaves. (a, c) Col-0, (b, d) *dxo1-2*, (a, b) leaves at the vein, (c, d) leaves structure. Red lines show the border between palisade and spongy mesophyll cells. AdE—adaxial epidermis, AbE—abaxial epidermis, PM—palisade mesophyll, SM—spongy mesophyll, VT—vascular tissue. Bars represent the mean of three independent biological replicates with error bars showing SD, $P < 0.05$ for Tukey's test.

**Fig. S2**

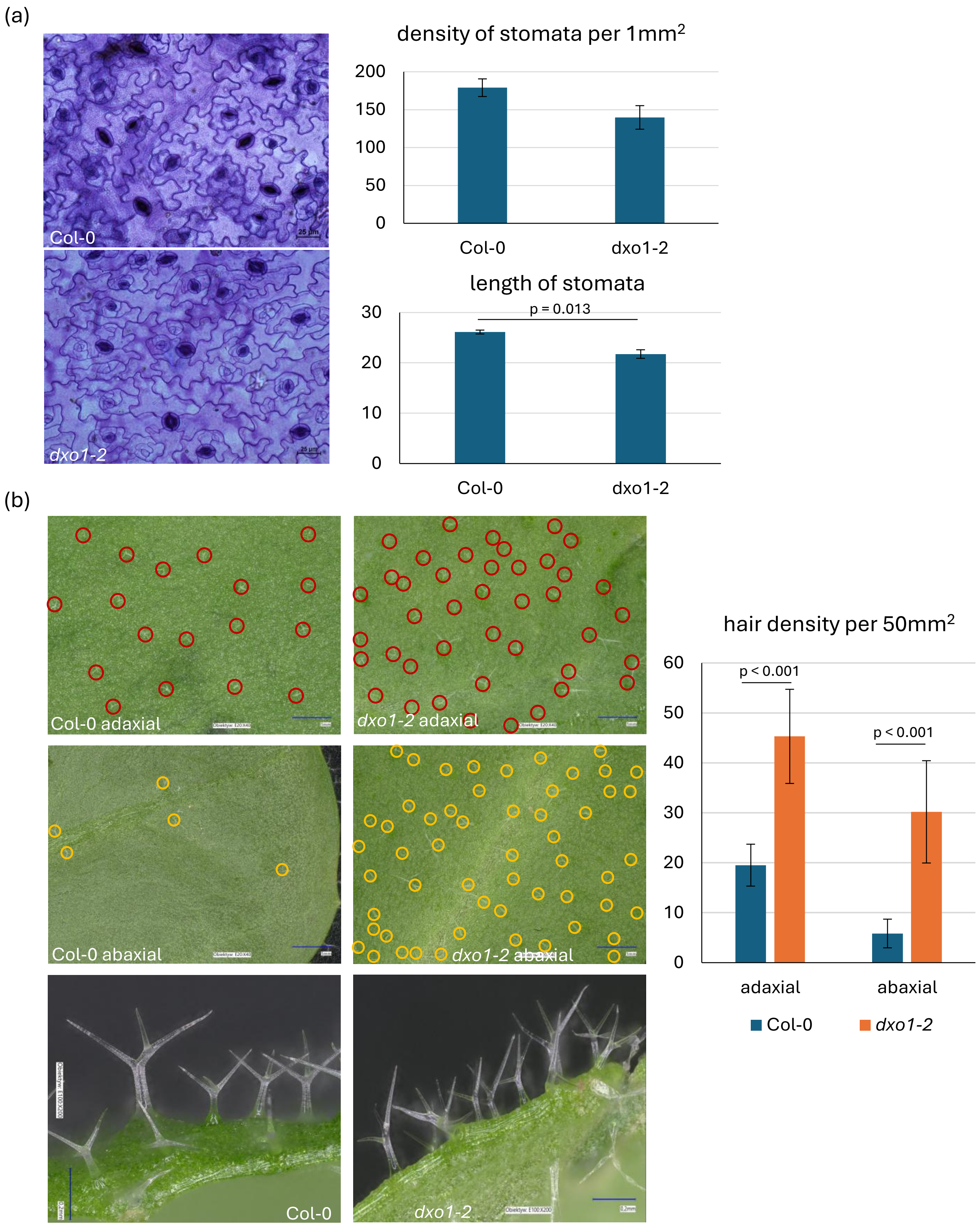


**Fig. S2** Leaf morphology in Col-0 and *dxo1-2* plants. (a) Stomata density and aperture length on the leaf surface of 6-week-old Col-0 and *dxo1-2* plants were estimated using a Nikon Eclipse 80i microscope (Nikon). Scale bar, 25 µm. Bars represent the mean of five independent biological replicates with error bars showing SD, $P < 0.05$ for Tukey's test. (b) Trichome density on the leaf surface of 6-week-old Col-0 and the *dxo1-2* plants. Images of the abaxial and adaxial leaf surfaces were captured using a digital microscope (VHX-7000N Keyence). Each trichome is circled in red (adaxial) or yellow (abaxial) to facilitate comparison of trichome density. Four leaves were analyzed for each genotype. Bars represent the mean of three independent biological replicates with error bars showing SD, $P < 0.05$ for Tukey's test.

**Fig. S3**

| Genes name | *dxo1-2* $MgCl_2$ vs Col-0 $MgCl_2$ | | *dxo1-2 Pst* vs *dxo1-2* $MgCl_2$ | | *dxo1-2 Pst* vs Col-0 *Pst* | | Col-0 *Pst* vs Col-0 $MgCl_2$ | |
|---|---|---|---|---|---|---|---|---|
| | 3'RNA-seq | northern blot | 3'RNA-seq | northern blot | 3'RNA-seq | northern blot | 3'RNA-seq | northern blot |
| *PR1* | 24.08 | 10.60 | 1.51 | 2.26 | 29.04 | 8.00 | 1.25 | 3.00 |
| *PR2* | 6.19 | 0.23 | 1.09 | 3.60 | 4.66 | 1.66 | 1.45 | 9.29 |
| *PR5* | 9.00 | 3.69 | 1.25 | 1.30 | 8.82 | 3.48 | 1.27 | 1.38 |
| *GSTF6* | 2.71 | 1.96 | 1.37 | 1.76 | 1.77 | 1.31 | 2.11 | 2.64 |
| *JAZ1* | 0.85 | 0.69 | 1.48 | 6.09 | 1.11 | 0.82 | 1.13 | 5.13 |
| *PDF1.2* | 0.26 | 0.29 | 0.99 | 1.00 | 0.02 | 0.05 | 14.32 | 5.14 |

**Fig. S3** Comparison of the expression level of selected pathogen response genes in Col-0 and the *dxo1-2* mutant based on RNA-seq and northern blot analysis. Numbers represent fold change.

**Fig. S4**

(a)

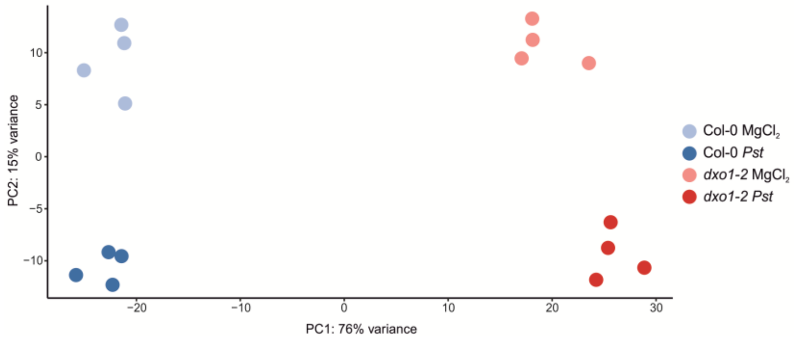


(b)

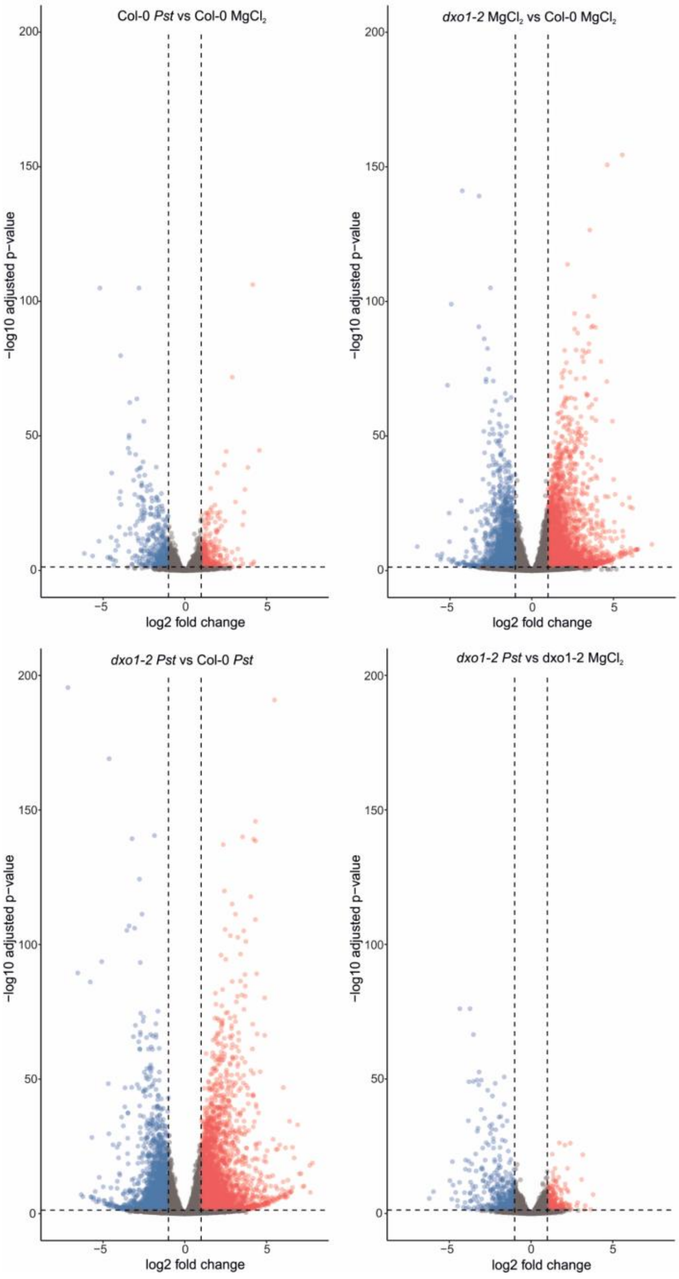


**Fig. S4** Results of RNA-seq analysis. (a) PCA (Principal component analysis) shows that biological replicas in 3′RNA-seq create four groups based on the presence of the *dxo1-2* mutation and *Pst* infection. (b) Volcano plots depicting the fold change in mRNA levels with comparisons between different conditions indicated above each plot. Significantly affected mRNAs (DESeq2; padj < 0.05) are represented by colours: upregulated (log2FC > 1 – red) and downregulated (log2FC < -1 – blue).
